# A DUF1068 protein acts as a pectin biosynthesis scaffold and maintains Golgi morphology and cell adhesion in Arabidopsis

**DOI:** 10.1101/2021.05.03.442108

**Authors:** Rahul S. Lathe, Heather E. McFarlane, Ghazanfar Abbas Khan, Berit Ebert, Eduardo Antonio Ramírez-Rodríguez, Niels Noord, Rishikesh Bhalerao, Staffan Persson

## Abstract

Adjacent plant cells are connected by specialized cell wall regions, called middle lamellae, which influence critical agricultural characteristics, including fruit ripening and organ abscission. Middle lamellae are enriched in pectin polysaccharides, specifically homogalacturonan (HG). Here, we identify a plant-specific Arabidopsis DUF1068 protein, called NKS1, that is required for middle lamellae integrity and cell adhesion. NKS1 localises to the Golgi apparatus and loss of the protein results in changes to Golgi structure and function. The *nks1* mutants also display HG deficient phenotypes, including reduced seedling growth, changes to cell wall composition, and tissue integrity defects. These phenotypes are identical to those of the HG deficient mutants *qua1* and *qua2*. Notably, NKS1 physically interacts with both QUA1 and QUA2, and genetic interaction analyses reveal that they work in the same pathway. Based on these results we propose that NKS1 works as a scaffold for HG synthesis and that such scaffolding is important to support Golgi function and the organization of the pectin synthesis machinery.

## INTRODUCTION

Growing plant cells are surrounded by a primary cell wall: a strong yet flexible extracellular matrix that is largely made of polysaccharides. Cell walls have the strength to resist turgor pressure and to direct cell morphology, but they are flexible enough to allow plant cells to expand. Long, strong cellulose microfibrils are the main load-bearing components of primary cell walls and are embedded in a hydrated matrix of pectins and hemicelluloses, with some proteins (Anderson & Kieber, 2020). Pectins are a heterogeneous class of acidic polysaccharides that are divided into Homogalacturonan (HG), Rhamnogalaturonan (RG) I and RGII (Atmodjo et al., 2013; Anderson, 2016). Pectins are particularly abundant in the primary cell walls of dicots, such as the model plant, *Arabidopsis thaliana*.

Pectins are made in the Golgi apparatus by the coordinated action of transporters and enzymes. Sugar interconversion enzymes, which are generally cytosolic, generate the nucleotide sugar building blocks for pectin synthesis (Temple et al., 2016); nucleotide sugar transporters facilitate their movement across Golgi membranes (Rautengarten et al., 2014); glycosyltransferases (GTs) catalyze their incorporation into pectic polysaccharides (Bouton et al., 2002); and methyltransferases and acetyltransferases further modify some pectins (Mouille et al., 2007). In particular, HG is secreted in a highly methylesterified form (Zhang & Staehelin, 1992). Once in the cell wall, HG may be modified by de-esterification, which can affect pectin crosslinking via Ca^2+^, and ultimately influence the mechanical properties of the cell wall (Peaucelle et al., 2011; Peaucelle et al., 2015). Indeed, a feedback loop exists between mechanical forces and pectin synthesis (Verger et al., 2018), and pectins are implicated in plant cell morphogenesis (Peaucelle et al., 2015; Bidhendi et al., 2019; Haas et al., 2020). For example, during growth symmetry breaking in the Arabidopsis hypocotyl, changes to pectin structure precede other changes in the cell cortex and cell wall, including cortical microtubule reorientation and realignment of cellulose deposition (Peaucelle et al., 2015).

Adjacent plant cells are connected by specialized regions of the cell wall, called middle lamellae. Regulation and degradation of middle lamellae underly critical agricultural characteristics, including fruit ripening (Uluisik et al., 2016) and organ abscission (Rhee et al., 2003; Ogawa et al., 2009), such as seed pod shattering (Lewis et al., 2006). Middle lamellae are pectin-rich and particularly enriched in HG (Willats et al., 2001). Therefore, defects in HG synthesis can lead to loss of cell-cell adhesion and epidermal tissue integrity, with dramatic consequences for plant growth and development (Bouton et al., 2002; Mouille et al., 2007). Such defects are evident in mutants that affect a member of the GT8 family of putative galacturonosyl transferases (GalATs) called QUASIMODO (QUA)1; *qua1* mutants had reduced levels of HG and displayed epidermal cell separation (Bouton et al., 2002). Similar phenotypes are observed in plants with mutations that affected a potential pectin methyltransferase, QUA2 (Mouille et al., 2007). While the pectin methyltransferase activity of QUA2 has not yet been demonstrated, a close homolog of QUA2, QUA3, was subsequently shown to harbor such activity (Miao et al., 2011).

While pectin synthesis occurs in the Golgi apparatus, there is contradictory data as to whether proteins required for HG synthesis are distributed across different Golgi cisterna (Zhang & Staehelin, 1992; Parsons et al., 2019), or whether HG synthesis proteins act as part of multiprotein complexes (Atmodjo et al., 2011; Harholt et al., 2012; Atmodjo et al., 2013), or a combination of both models (Zabotina et al., 2021; Hoffmann et al., 2021). A better understanding of the pectin synthesis machinery and its interactors is required to appreciate the synthesis of this class of polysaccharides and to open potential for pectin engineering for agricultural improvements. Here, we report that a plant-specific Golgi-localized protein of unknown function (DUF1068) interacts with QUA1 and QUA2 to support HG synthesis, Golgi integrity and cell adhesion. We propose that this DUF1068 protein is a scaffold for HG synthesis and that such scaffolding is important to support Golgi function and the organization of the pectin synthesis machinery.

## RESULTS

### A DUF1068 protein, referred to as NKS1, is required for cell elongation

Co-expression is a powerful approach to identify functionally related genes (Usadel et al., 2009). Using ATTED-II (Obayashi et al. 2018), we identified several genes from the Arabidopsis DUF1068 family as co-expressed with primary wall *CELLULOSE SYNTHASE* (*CESA*) genes and pectin biosynthesis-related genes, including *GALACTURONOSYL TRANSFERASES* (GalATs) *9* (*GAUT9*) and *QUA1*, the HG methyltransferase *QUA3*, and many S-adenosylmethionine family transporter genes, which might play important roles in HG-methyltransferase activity (Table S1 and S2). We tested T-DNA lines that were annotated to target DUF1068 genes from the co-expression list above. Of the ones we tested, two independent T-DNA lines targeting the DUF1068 gene *At4g30996* (also called *Na^+^ AND K^+^-SENSITIVE 1* (*NKS1*); Choi et al., 2011) displayed significant reduction in mean hypocotyl length of six-day-old etiolated seedlings, compared to wild type (Figures 1B, 1C). Moreover, growth kinematics of etiolated seedlings were dramatically affected in the *nks1* mutant hypocotyls compared to wild type (Figure 1D). We refer to these two T-DNA lines as *nks1-1* (SALK_15107) and *nks1-2* (GK-228H05) (Figure S1A). Whereas RT-PCR analysis indicated that the two lines were transcriptional null lines (Figure S1B), qPCR analyses revealed some residual *NKS1* expression in *nks1-1* (Figure 1A). Nevertheless, there was a substantial reduction in *NKS1* expression in the two T-DNA lines and the growth phenotypes of *nks1-1* and *nks1-2* seedlings could be rescued by molecular complementation using fluorescent protein fusions to NKS1, either NKS1-GFP or GFP-NKS1 (Figure 1E). Although NKS1 is ubiquitiously expressed, we primarily obeserved phenotypes in young seedlings (Figure S1C).

**Figure 1:**
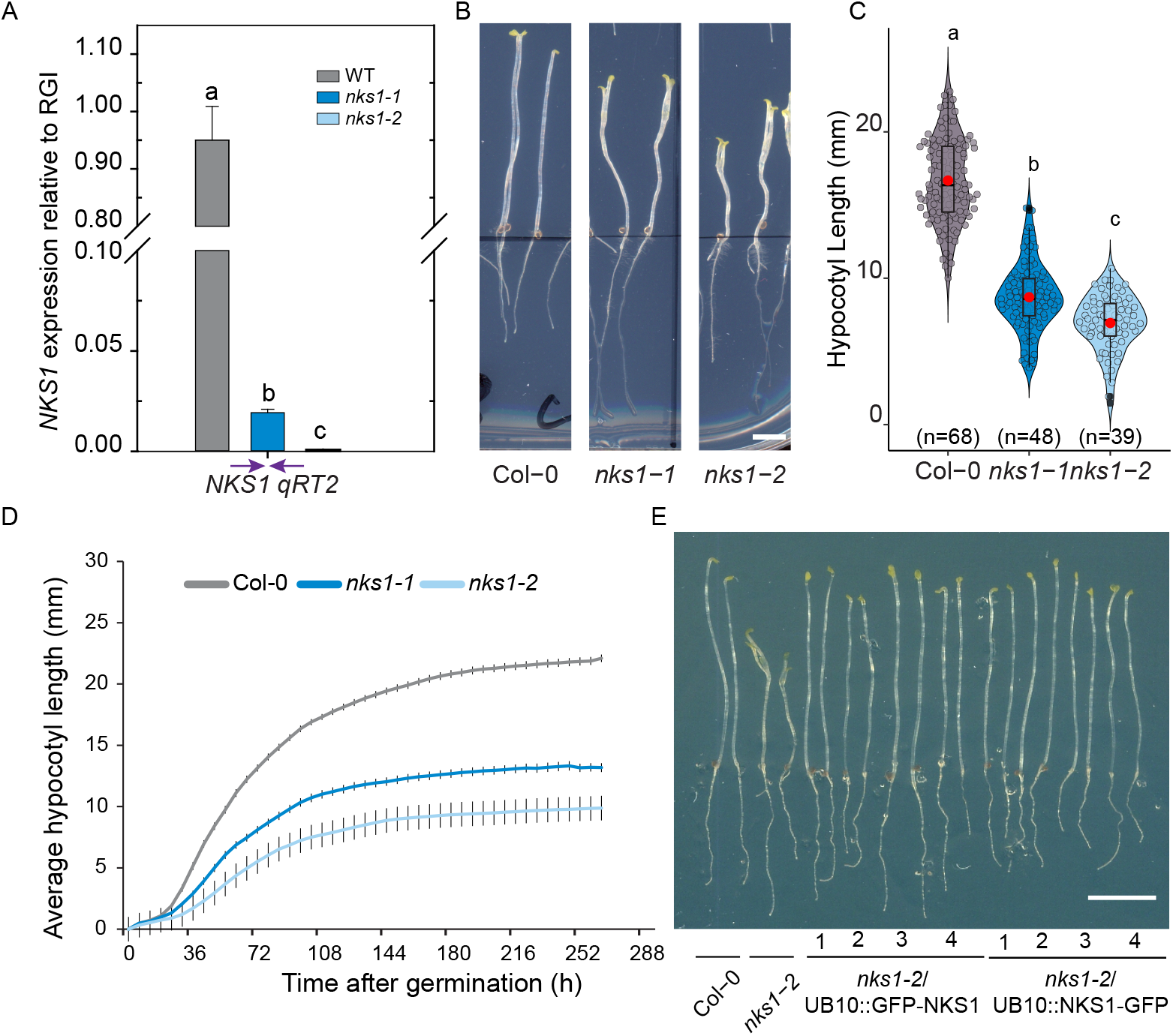
*nks1* mutants are defective in cell elongation. (A) Quantitative reverse-transcription polymerase chain reaction (qRT-PCR) of *NKS1* transcript levels normalized to relative gene index (RGI) from Col-0, *nks1-1* and *nks1-2*; bars represent means of three biological replicates ±SD. (B) Representative images of six-day old etiolated seedlings of Col-0, *nks1-1* and *nks1-2*. (C) Quantification of hypocotyl lengths from six-day-old etiolated seedlings of Col-0, *nks1-1* and *nks1-2*; data distribution is outlined by the shape, plot box limits indicate 25^th^ and 75^th^ percentiles, whiskers extend to 1.5 times the interquartile range, median is indicated by a horizontal line, mean by a red dot, individual data points are shown, and n (seedlings) is indicated in parentheses. (D) Etiolated hypocotyl growth kinematics of Col-0, *nks1-1* and *nks1-2* seedlings (n=15 seedlings); points indicate mean ±SD). (E) Representative images of pUB10-GFP-NKS1 and pUB10-NKS1-GFP expressed in the *nks1-2* background along with controls (Col-0 and *nks1-2*). Letters in (A) and (C) specify statistically significant differences among samples as determined by one way ANOVA followed by Tukey’s HSD test (p < 0.05). Scale bars represent 2 mm in (B) and 5 mm in (E).

### Functional fluorescently-tagged NKS1 fusions localize to the *medial*-Golgi apparatus

To better understand NKS1 function, we undertook subcellular localization studies of the functional NKS1-GFP and GFP-NKS1 lines (Figure 1E). Both NKS1-GFP and GFP-NKS1 were localized to doughnut shaped particles that were rapidly streaming in the cytoplasm of hypocotyl epidermal cells (Figure 2A), which is typical of Golgi-or trans-Golgi Network (TGN) localized proteins. We also generated a NKS1-mRFP fusion for colocalization purposes that displayed similar localization to both GFP fusions. Quantitative colocalization with markers for the ER (HDEL; Batoko et al., 2000), the Golgi apparatus (WAVE18/Got1P homolog and WAVE22/SYP32, Geldner et al., 2009), the *trans*-Golgi Network (TGN; VHAa1, Dettmer et al., 2006), and late endosomes (WAVE2/RabF2b and WAVE7/RabF2a, Geldner et al., 2009) revealed that NKS1-GFP co-localized with Golgi markers and displayed some overlap with TGN markers (Figure 2B; Figure S2A). To distinguish between the Golgi and TGN, we treated seedlings with Brefeldin A (BFA) for 60 minutes, which in Arabidopsis root cells causes aggregation of TGN and other compartments into BFA bodies, while intact Golgi stacks surround the core of the BFA body (Geldner et al., 2003; Grebe et al., 2003; Gendre et al., 2011). After BFA treatment, NKS1-GFP localized to discrete puncta around the core of the BFA body, and NKS1-GFP remained highly colocalized with the Golgi marker, XYLT (Saint-Jore-Dupas et al., 2006) but was no longer colocalized with the TGN marker, VHAa1, which was in the core of the BFA bodies (Dettmer et al., 2006) (Figure S2B). These data are consistent with those of subcellular proteomics studies, which have detected NKS1 in Golgi fractions (Nikolovski et al., 2012; Parsons et al., 2012; Parsons et al., 2019).

**Figure 2.**
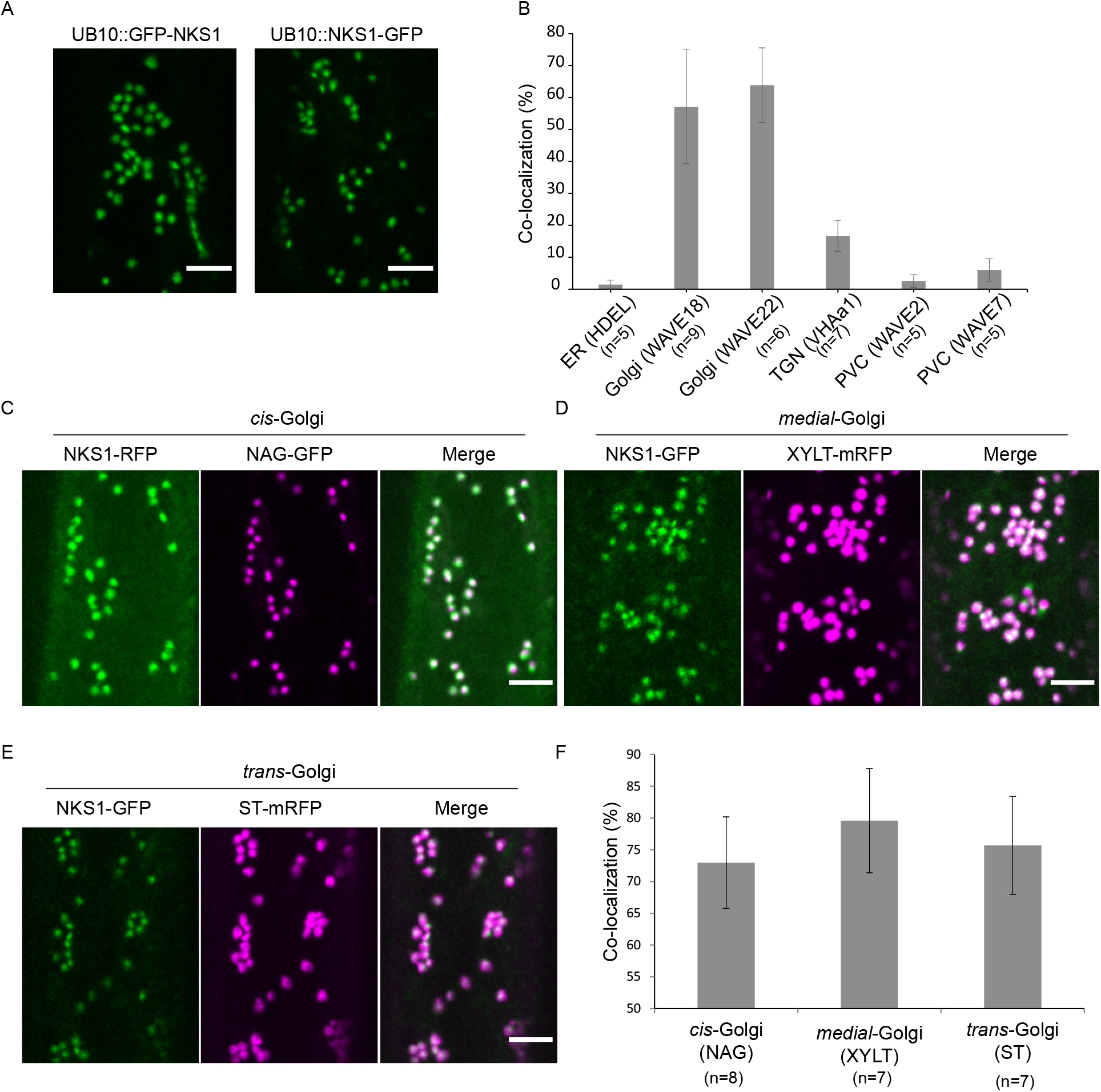
Functional NKS1-GFP fusion is localized to the *medial*-Golgi apparatus. (A) Representative images NKS1 localization to endomembrane compartments; both N- and C-terminal GFP fusion construct localization in single focal plane images of hypocotyl epidermal cells of 3-day-old etiolated seedlings. (B) Quantification of co-localization between NKS1 and various endomembrane compartments specific markers. (C), (D) and (E) Representative images of colocalization between NKS1-GFP and Golgi cisternae markers: NAG-GFP (*cis*-Golgi), XYLT-mRFP (*medial*-Golgi) and ST-mRFP (*trans*-Golgi) in hypocotyl epidermal cells of 3-day-old etiolated seedlings. (F) Quantification of colocalization percentage between NKS1 and Golgi-cisternae specific markers. In bar charts, bars represent mean ±SD, n (cells, one cell imaged per seedling) is indicated in parentheses. Scale bars represent 5 μm in (A), (C), (D) and (E).

Different Golgi cisternae are associated with different biochemical functions, *i.e*., the assembly or modification of certain cell wall components (Zabotina et al., 2021; Hoffmann et al., 2021). To investigate whether NKS1 is associated with certain cisternae, we next crossed the NKS1-GFP or NKS1-RFP fluorescent lines with markers for the *cis*-Golgi (NAG, Grebe et al., 2003), *medial*-Golgi (XYLT, Saint-Jore-Dupas et al., 2006) or *trans*-Golgi (ST, Renna et al., 2005) to generate dual-labelled fluorescent lines. While NKS1 co-localized with all three markers, the highest degree of colocalization was observed with *medial*-Golgi markers (Figure 2C–2F). Together, these results confirm that the functional NKS1-GFP fusion is preferentially localized to medial-cisternae of the Golgi apparatus.

### NKS1 is a plant-specific transmembrane protein with its DUF1068 domain inside the Golgi lumen

NKS1 encodes a plant-specific protein of 172 amino acids with a predicted molecular mass of 19 KDa. Genes encoding DUF1068-containing proteins are found throughout land plants (Embryophyta), including *Marchantia polymorpha* and *Physcomitrium patens*, suggesting that DUF1068 function was acquired as plants colonized land (Figure S3A). NKS1 is predicted to contain one transmembrane domain (TM; TmHMM server, Krohg et al., 2001) (Figure S3B). This prediction also suggested that the first 17 amino acids in the N-terminus of NKS1 are cytoplasmic, which would imply that the amino acids after the TM domain would face the Golgi lumen. To test this prediction, we used a GO-PROMPTO assay (Søgaard et al., 2012). Here, we fused the N-terminal part of VENUS (Vn; the first 155 amino acids), or the C-terminal part of VENUS (Vc; the last 84 amino acids), in frame either before (cytosolic reporter) or after (Golgi luminal reporter) the first 52 amino acids of the rat ST protein (TMD), which consists of a transmembrane domain targeted to the Golgi apparatus. We observed clear fluorescence complementation only when co-expressing Vc-NKS1 with the cytosolic reporter, but not with the luminal reporter (Figure S3C). These results corroborate that the N-terminus of NKS1 faces the cytoplasm, while the bulk of the protein, including the DUF1068 domain, is in the Golgi lumen (Figure S3D).

### *nks1* mutants are defective in Golgi structure and function

The Golgi localization of NKS1 prompted us to examine the structure and function of the Golgi apparatus in *nks1* mutants. We therefore generated double Golgi marker lines that carried the *cis*-Golgi marker NAG-EGFP (Grebe et al., 2003) and the *trans*-Golgi marker ST-mRFP (Renna et al., 2005). Simultaneous dual colour live cell imaging and object-based colocalization between the two markers demonstrated that these two Golgi markers were significantly further apart in *nks1-2* mutants, relative to wild type (Figure 3A–3C). This increased separation between *cis*-Golgi and *trans*-Golgi in *nks1* mutants was not an artefact of faster Golgi stack movement within cells; in fact, measurements of Golgi marker dynamics indicated that Golgi stacks moved significantly slower in *nks1-2* mutants, relative to wild type (Figure 3D). We therefore examined Golgi structure at high resolution using transmission electron microscopy (TEM) of high-pressure frozen, freeze-substituted hypocotyls and found that Golgi morphology was dramatically affected in *nks1* mutants (Figure 3E). We frequently observed curved Golgi stacks in both alleles of *nks1*, and the proportion of curved Golgi stacks was significantly higher in *nks1* mutants than wild type (Figure 3F). Loss of NKS1 also resulted in fewer cisternae per Golgi stack but no changes to other Golgi morphometrics (cisternal length:width, proportion of Golgi stacks with an associated TGN) (Table S5). Dual-axis transmission electron tomograms of wild type and *nks1-2* Golgi stacks confirmed that Golgi curving was not an artefact of the plane of section and provided additional insight into the Golgi structure defects observed in *nks1* mutants (Figure 3G).

**Figure 3:**
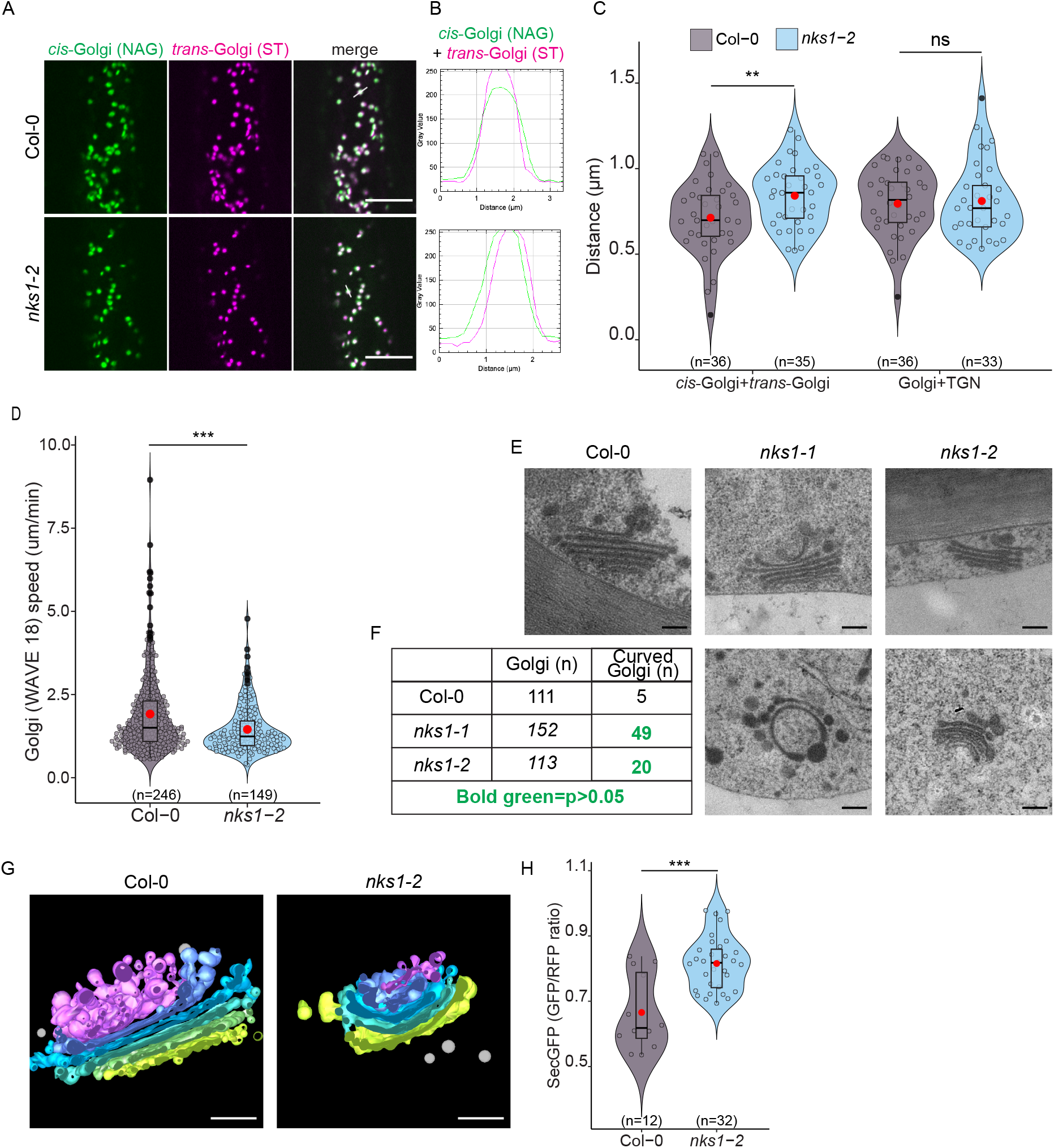
*nsk1* mutants are defective in Golgi apparatus structure and function. (A) Representative images of simultaneous dual wavelength localization of *cis*-Golgi (NAG) and *trans*-Golgi (ST) dual markers in Col-0 and *nks1-2* hypocotyl epidermal cells of 3-day-old etiolated seedlings. (B) Linescan graph showing distance between *cis*-Golgi (NAG) and *trans*-Golgi (ST) dual markers in Col-0 and *nks1-2* from single Golgi particle shown in A. (C) Quantification of the distance between *cis*-Golgi (NAG) and *trans*-Golgi (ST) dual markers or medial-Golgi (WAVE18) and TGN (VHAa1) dual markers in Col-0 and *nks1-2* hypocotyl epidermal cells of 3-day-old etiolated seedlings. (D) Quantification of Golgi (WAVE 18) speed in Col-0 and *nks1-2* cells. (E) Representative transmission electron microscopy images of Golgi ultrastructure from Col-0, *nks1-1* and *nks1-2* hypocotyl epidermal cells of 3-day-old etiolated seedlings. (F) Quantification of the frequency of Golgi curving in Col-0 and *nks1* alleles. Statistically significant numbers are shown in bold green color (p<0.05, χ^2^ test, 1 d.f.). (G) Representative electron tomogram models of Col-0 and *nks1-2* Golgi apparatus; the *cis*-most cisterna is labelled in yellow, the *trans*-most cisterna in purple, and cisternae between are labelled by a gradient of green through blue, the TGN is labelled in pink and free vesicles in in grey. (H) Quantification of SecGFP secretion ratio in Col-0 and *nks1-2* hypocotyl epidermal cells of 3-day-old etiolated seedlings. Asterisks in (C), (D) and (H) indicate statistically significant difference between Col-0 and *nks1* as determined by unequal variance, two tailed Student’s t test, where ***p < 0.0005, **p < 0.005. In violin plots, data distribution is outlined by the shape, plot box limits indicate 25^th^ and 75^th^ percentiles, whiskers extend to 1.5 times the interquartile range, median is indicated by a horizontal line, mean by a red dot and individual data points are shown, and n is indicated in parentheses. Scale bars represent 10 μm in (A), and 200 nm in (E) and (G).

To determine whether the structural changes to the Golgi apparatus affected Golgi function in *nks1* mutants, we assayed a ratiometric marker of soluble protein secretion, sec-GFP (Samalova et al., 2006). Sec-GFP is GFP fused to a signal peptide, which directs the protein to the secretory pathway and ultimately to the apoplast, where the GFP fluorescence is quenched by the low pH; because of the stochastic expression of sec-GFP, especially in epidermal cells, an endomembrane-targeted RFP is produced in equal amounts to sec-GFP; therefore, the ratio of GFP:RFP can be compared across different plants (Samalova et al., 2006). The ratio of GFP:RFP was significantly higher in *nks1-2* mutants compared to wild type (Figure 3H; Figure S4A), indicating a secretion defect.

Since secretion flows through both the Golgi apparatus and the TGN, we tested whether TGN structure or function was affected in *nks1* mutants. Using simultaneous dual colour live cell imaging and object-based colocalization, we found no significant difference in the distance between a Golgi marker (WAVE18, Geldner et al., 2009) and TGN marker (VHAa1, Dettmer et al., 2006) between wild type and *nsk1* (Figure 3C; Figure S4B). There were also no substantial differences in Golgi-TGN association or TGN morphology at the TEM level (Figure S4C). To examine anterograde trafficking from the TGN, we tracked the localization of PIN2-GFP (Xu & Scheres, 2005) in response to BFA. Since BFA-treatment of Arabidopsis root epidermal cells induces aggregation of TGN and endosomes in the BFA body, but leaves Golgi stacks intact and clustered around the BFA body (Geldner et al., 2003; Grebe et al., 2003; Gendre et al., 2011), signal recovery after BFA washout primarily involves protein secretion from the BFA body/TGN to the plasma membrane. We found no significant differences between the ratio of PIN2-GFP plasma membrane signal compared to intracellular signal or in the number of BFA bodies between wild type and *nks1-2* mutants at any stage of BFA treatment or washout (Figure S4D). Finally, since the plant TGN also functions as an early endosome (Viotti et al., 2010), we assayed endocytosis by tracking uptake of the fluorescent endocytic marker, FM4-64 (Bolte et al., 2004). There were no significant differences in FM4-64 uptake between wild type and *nks1-2* (Figure S4E). Together, these results indicate that while TGN structure and function seem unaffected by loss of NKS1, Golgi apparatus structure and function are impaired in *nks1* mutants.

### *nks1* mutants are defective in cell adhesion and cell wall pectins

In addition to the defects in cell elongation, *nks1-1* and *nks1-2* mutants displayed defects in cell adhesion: in cryo-scanning electron microscopy (cryo-SEM), hypocotyl cells of *nks1* mutants seemed to be peeling apart in both epidermal and cortical cell layers (Figure 4A & 4B). Consistent with a loss of tissue integrity, *nks1* mutant hypocotyls were permeable to toluidine blue dye (Figure S5A).

**Figure 4:**
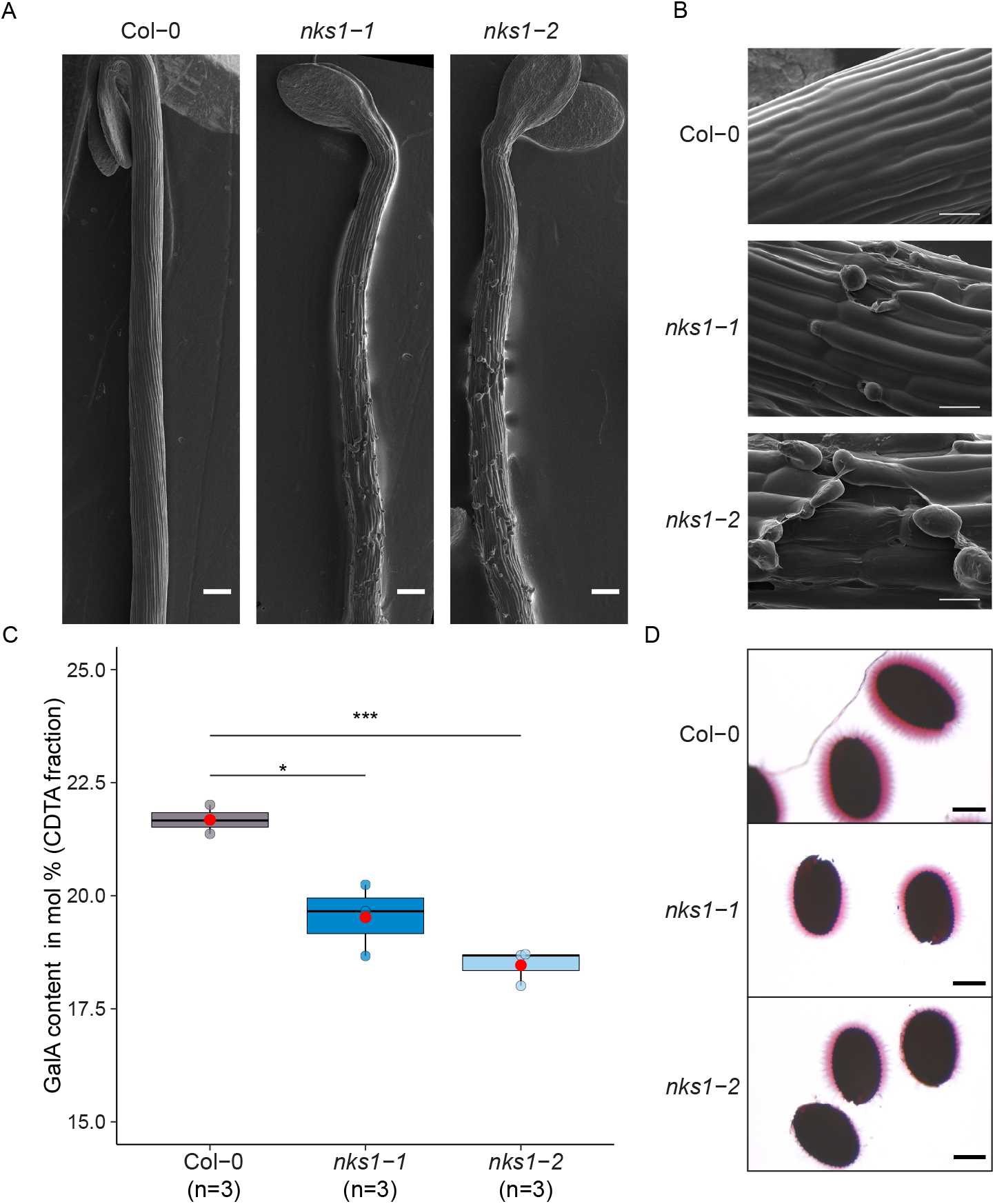
*nks1* mutants are defective in cell adhesion and cell wall pectins. (A) Representative scanning electron microscopy of 5-day old etiolated seedlings of Col-0, *nks1-1* and *nks1-2*. (B) Higher magnification of the seedlings shown in (A) showing epidermal cell layer in Col-0 and *nks1* alleles. (C) GalA levels in Col-0, *nks1-1* and *nks1-2* in the CDTA-extracted cell wall fraction as measured by HPAEC-PAD. (D) Seed Mucilage staining of Col-0, *nks1-1* and *nks1-2* with Ruthenium Red solution. Asterisks in (C) indicate statistically significant difference between Col-0 and *nks1-2* as determined by unequal variance, two tailed Student’s t test, where ***p < 0.0005, *p < 0.05. Data is shown in boxplot where plot box limits indicate 25^th^ and 75^th^ percentiles, whiskers extend to 1.5 times the interquartile range, median is indicated by a horizontal line, mean by a red dot and individual data points are shown, and n (distinct pools of homogenized seedlings) is indicated in parentheses. Scale bars represent 200μm in (A), 50μm in (B), 200μm in (D).

The cell walls of adjacent plant cells are joined by the middle lamella, a pectin-rich region that is particularly enriched in HG (Willats et al., 2001) and changes in cell wall HG can therefore lead to cell-cell adhesion defects and loss of epidermal tissue integrity (Bouton et al., 2002; Mouille et al., 2007). HG and other pectins are characterized by high levels of galacturonic acid (GalA) (Atmodjo et al., 2013). Therefore, we quantified total cell wall monosaccharides by HPAEC-PAD. These experiments revealed a significant reduction in GalA content compared to wild type in *nks1-2*, which was accompanied by a significant increase in arabinose content compared to wild type (Figure 4C; Table S3). Sequential extraction of cell wall polymers confirmed that a significant decrease in GalA in both *nks1* alleles was associated with the CDTA-extracted fraction that mainly extracts calcium cross-linked pectins from the cell wall. *nks1* mutants also displayed other pectin defective phenotypes, including reduced seed coat mucilage (Western et al., 2000) (Figure 4D). Despite *NKS1* coexpression with primary wall *CESA* genes (Table S1), we did not observe any significant differences in cellulose content between *nks1-2* and wild type seedlings (Figure S5B). Similarly, there were no significant changes in fluorescently-tagged CESA dynamics in the plasma membrane (Paredez et al., 2006) in *nks1-2* mutant hypocotyl cells, compred to wild type (Figure S5C).

### *nks1* mutants phenocopy *qua1* and *qua2* pectin synthesis mutants and NKS1 interacts with QUA1 and QUA2

The cell wall pectin and cell adhesion defects of *nks1* mutants were reminiscent of *qua1* (Bouton et al., 2002) and *qua2* mutants (Mouille et al., 2007), and *NKS1* was tightly co-expressed with *QUA1* and *QUA3* (Table S1). QUA1 is similar to GT8 family GalATs and QUA2 is putative methyltransferase; both have been implicated in HG synthesis (Bouton et al., 2002; Mouille et al., 2007). We therefore investigated whether *nks1* mutants shared other physiological, molecular, and genetic phenotypes with *qua1* and *qua2* mutants.

Cell adhesion mutants, including *qua1* and *qua2*, display increased pectin related cell wall integrity signaling (Verger et al., 2016), such as increased expression of *FAD-LINKED OXIDOREDUCTASE* (*FADLox*), a marker gene associated with pectin responses (Denoux et al., 2008; Kohorn et al., 2014). Similar to that of the *qua* mutants, *nks1-1* and *nks1-2* showed significant increase in *FADLox* expression compared to wild type (Figure S6A). The *nks1* mutants also displayed increased accumulation of anthocyanins when grown on high sucrose containing growth media (Figure S6B), which was observed in the *qua1-1* and *qua2-1* mutants (Verger et al., 2016, Bouton et al., 2002, Gao et al., 2008; Krupkova et al., 2007).

Recently, Verger et al., 2018 documented the importance of epidermal continuity for mechano-perception. By modulating turgor (by changing the osmotic potential of the growth media) they could rescue cell-adhesion defects in *qua1* and *qua2* mutants, possibly through a tension-adhesion mechanism connected to cortical microtubules (Verger et al., 2018). To test whether we also could restore the cell adhesion defects in *nks1* mutants, we grew seedlings on media with reduced osmotic potential, *i.e*., on “hard” media (2.5% agar; Verger et al., 2018) compared to control (0.8% agar). Interestingly, cell elongation and cell adhesion defects were significantly restored when *nks1* seedlings were grown on the hard media (Figure S6C).

Mutations in *ESMD1*, which encodes a putative O-fucosyltransferase GT106 family protein, suppress the *qua1-1* and *qua2-1* growth and cell adhesion phenotypes. Introducing *esmd1-1* into *nks1-2* also suppressed the hypocotyl elongation and cell adhesion phenotypes of *nks1-2* (Figure 5A & 5B), implying that loss of *NKS1*, *QUA1*, and *QUA2* all affect the same cell wall sensing and/or response pathway. To directly test this hypothesis, we generated double mutants between *nsk1-2* and *qua2-1*. Because the *qua1-1* is in the Ws-4 background, we focused our efforts on the *qua2-1* which, like the *nks1* alleles, is in a Col-0 background. We found that *nsk1-2 qua2-1* double mutants resembled the single mutants, which is consistent with the hypothesis that *NKS1* and *QUA2* act in the same complex or pathway (Figure 5C & 5D).

**Figure 5:**
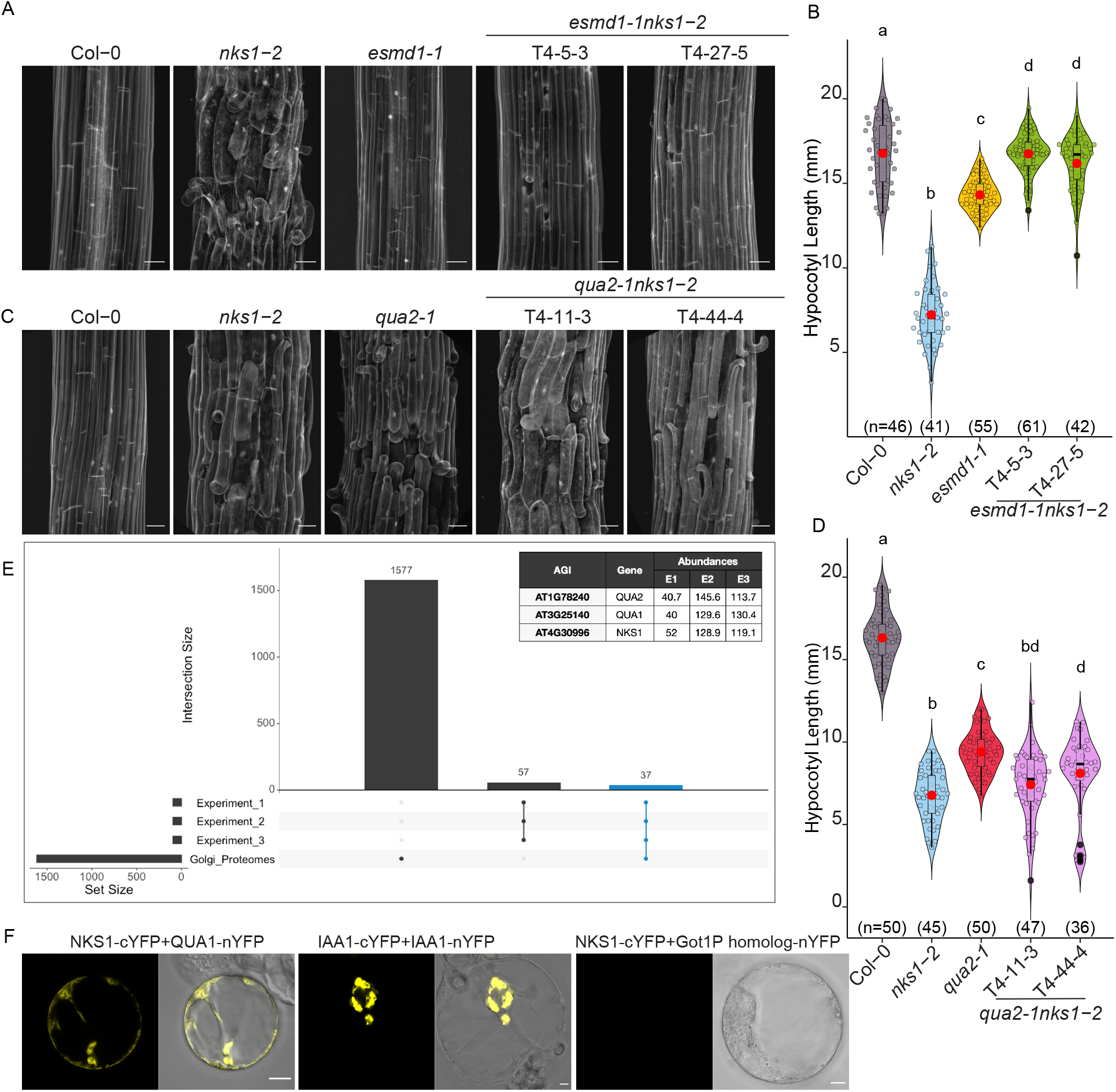
NKS1 interacts with QUA1 and QUA2. (A) Representative z-projections (sum averages) of confocal stacks from propidium iodide-stained etiolated five-day old hypocotyl epidermal cell files from Col-0, *nks1-2*, *esmd1-1* and two independent lines of *esmd1-1 nks1-2* double mutants. (B) Quantification of hypocotyl lengths of six-day old etiolated seedlings from Col-0, *nks1-2*, *esmd1-1* and *esmd1-1 nks1-2* double mutants. (C) Representative z-projections (sum averages) of confocal stacks from propidium iodide-stained etiolated five-day old hypocotyl epidermal cell files from Col-0, *nks1-2*, *qua2-1*, and two independent lines of *qua2-1nks1-2* double mutants. (D) Quantification of hypocotyl lengths of six-day old etiolated hypocotyls from Col-0, *nks1-2*, *qua2-1* and *qua2-1 nks1-2* double mutants. (E) UpSet Plot comparing proteins identified in three independent NSK1-GFP immunoprecipitation experiments, compared to previously published Golgi proteomes, where total set size is indicated at the lower left and intersection set sizes (with intersections defined by joined dots) are indicated in the upper bar chart; blue set indicates 37 Golgi-localized proteins identified in all three experiments and in at least 6/8 biological replicates. Insert indicates relative abundance of NKS1, QUA1, and QUA2 in each independent experiment (n=8, two or three samples each in three independent experiments). (F) Representative images of bimolecular fluorescence complementation (BiFC) assay in Arabidopsis root cell culture protoplasts showing interaction between NKS1 (-cYFP) and QUA1 (GAUT8) (-nYFP). IAA1-cYFP and IAA1-nYFP was used as a positive control and NKS1 (-cYFP) and Got1P homolog (-nYFP) as negative control. Letters in (B) and (D) specify statistically significant differences among samples as determined by one way ANOVA followed by Tukey’s HSD test (p < 0.05). In violin plots, data distribution is outlined by the shape, plot box limits indicate 25^th^ and 75^th^ percentiles, whiskers extend to 1.5 times the interquartile range, median is indicated by a horizontal line, mean by a red dot and individual data points are shown, and n (seedlings) is indicated in parentheses. Scale bars represent 5μm in (A) and (C); 10μm in (F).

As the bulk of the NKS1 resides inside the Golgi lumen (Figure S3D), but the DUF1068 sequence does not harbour any hallmarks of enzymatic activity, we wondered whether NKS1 might physically interact with QUA1 and QUA2, potentially acting as a pectin synthesis scaffold. To test this hypothesis, we performed immunoprecipitation of NKS1-GFP followed by LC-MS/MS analysis. We identified 248 proteins that were present with ‘high’ confidence in all three experiments and in at least six out of the eight biological replicates. To further refine this list, we used SUBA (Hooper et al., 2017) to filter for proteins that are predicted to localize to the Golgi apparatus, resulting in 94 candidates (Figure 5E). Comparison of these results with previously published Golgi proteomes (Nikolovski et al., 2012; Parsons et al., 2012; Parsons et al., 2019) revealed that 40% of the proteins identified via NKS1-GFP immunoprecipitation were a subset of these Golgi proteomes (Figure 5E; Table S4). Importantly, QUA1 QUA2 were identified among the top putative NKS1 interactors in all three experiments and all eight biological replicates of NKS1-GFP immunoprecipitation (Table S4). To corroborate that NKS1 interacts with QUA1 and QUA2, we undertook Bimolecular Fluorescence Complementation (BiFC) assays using Arabidopsis root protoplasts. Here, we detected clear positive interactions between NKS1 and QUA1, which localized to small intracellular puncta, but we did not observe any signs of interaction between NKS1 and another Golgi localized protein, Got1p (Figure 5F).

Taken together, the similar physiological, molecular, and genetic phenotypes imply that NKS1, QUA1, and QUA2 act in the same pathway, which we confirmed by documenting their physical interaction in the Golgi apparatus.

## DISCUSSION

Domain of Unknown Function proteins are classified by sequence similarity to each other but not to any protein of known function and make up almost 22% of all proteins in the Pfam database (El Gebali et al., 2019). NKS1 belongs to the DUF1068 family, members of which are only found in land plants (Embryophyta), and almost all annotated DUF1068 proteins consist entirely of only the DUF1068 domain, making it difficult to deduce their function from protein sequence. Previous studies had implicated NKS1 in salt tolerance (Choi et al., 2011); we hypothesize that the high concentration of sucrose in the media used by Choi et al. (2011) exacerbated the cell wall phenotype, since there are complex relationships between sugar availability and cell wall integrity responses (Hamann et al., 2009; Englesdorf et al., 2018). Here we show that NKS1 maintains Golgi apparatus structure and function, and may act as a scaffold for pectin synthesizing proteins.

Changes in pectin synthesis have been correlated with changes to Golgi structure (Young et al., 2008; Wang et al., 2017). For example, in seed coat epidermal cells, which synthesize an extraordinary volume of pectic mucilage during their development, Golgi stacks showed swollen margins, many associated vesicles, and a complex *trans*-Golgi network, while these changes were not observed in mutants lacking a key pectin synthesis gene (Young et al., 2008). Whether these structural changes to the Golgi reflect an active remodeling of the endomembrane system or are a passive consequence of polysaccharide flux through the Golgi remains to be determined (Hoffmann et al., 2021). Notably, in mammalian (HeLa) cells, changes to Golgi protein interactions were correlated with loss of GT function and dramatic changes to Golgi structure (van Galen et al., 2014), implying an important relationship between Golgi structure and function. These data are consistent with our characterization of *nks1* mutants, in which Golgi structure and function were defective. While the relationship between Golgi structure and function remains elusive, modelling has demonstrated that both changes to Golgi lipid composition and changes to curvature-generating proteins (*i.e*., vesicle trafficking machinery) can influence Golgi shape (Campelo et al 2017). According to this model, changes to pectin synthesis in *nks1* Golgi stacks might passively reshape the Golgi apparatus due to changes in vesicle trafficking.

The phenotypes of *nks1* mutants are strikingly similar to *qua1* and *qua2* mutants, including reduced cell elongation, cell adhesion defects, and suppression of the phenotypes under hyperosmotic conditions or by loss of ESMD (Verger et al., 2016). QUA1 is a predicted GalAT implicated in HG synthesis (Bouton et al., 2002), while QUA2 is a putative HG methyltransferase (Mouille et al., 2007). NKS1 lacks any sequence features that might suggest it is directly involved in pectin synthesis. However, the interactions between NKS1 and QUA1 and QUA2 led us to hypothesize that NKS1 could play a role in organizing the pectin synthesis machinery in the Golgi apparatus by mediating close associations between QUA1 and QUA2. Previous studies of HG synthesis have documented interactions between GAUT1 and GAUT7 (Atmodjo et al., 2011). While enzymatic activity has only been documented for GAUT1 (Sterling et al., 2006), GAUT7 is required for proper GAUT1 localisation to the Golgi (Atmodjo et al., 2011), and GAUT7 can increase GAUT1 activity *in vitro* (Amos et al., 2018). HG is secreted in a highly methylesterified form, presumably to prevent it from forming calcium bridge-mediated aggregations before its incorporation into the cell wall. Quantitative immunolabelling of HG in pectin synthesizing Golgi stacks predicted that HG methylesterification is highly efficient and nearly simultaneous with HG backbone synthesis, suggesting that the enzymes for backbone formation and methylesterification must act in concert (Zhang & Staehelin, 1992). Therefore, we propose a model in which NKS1 mediates interactions between the putative GalAT, QUA1, and the putative HG methyltransferase, QUA2, thus acting as a scaffold for these proteins to facilitate efficient and coordinated HG synthesis and methylesterification before pectin secretion.

## ACKNOWLEDGEMENTS

Some live cell imaging was conducted using instruments that are part of the Biological Optical Microscopy Platform (BOMP) at University of Melbourne and electron microscopy was conducted using instruments that are part of the Melbourne Advanced Microscopy Facility. We thank Shuai Nie and Ching-Seng Ang for processing samples at the Melbourne Mass Spectrometry and Proteomics Facility of The Bio21 Molecular Science and Biotechnology Institute. R.S.L acknowledges PhD scholarship from Deutscher Akademischer Austausch Dienst (DAAD., PKZ:91540412 (formerly A/10/75281)) at MPIMP; Germany and postdoctoral grant from Kempe foundation (# SMK-1759) to RPB at UPSC, Sweden. H.E.M. acknowledges an ARC Discovery Early Career Researcher Award (DE170100054), NSERC Discovery Grant (2020-05959) and funding from the Canada Research Chairs program as Canada Research Chair in Plant Cell Biology. B.E. acknowledges ARC Future Fellowship and Discovery Project Awards (FT160100276, DP180102630) and ongoing support from the University of Melbourne Botany Foundation. E.A.R.R. is supported by a CONACyT Beca de Posgrado en el Extranjero (2020-000000-01EXTF-00193). G.A.K. acknowledges an ARC Discovery Early Career Researcher Award (DE210101200) and a Swiss National Science Foundation Grant (P400PB_180834/1). N.N acknowledges funding from Erasmus+ (NL GRONING03). S.P. acknowledges the financial aid of an ARC Discovery grant (DP19001941), Villum Investigator (Project ID: 25915), DNRF Chair (DNRF155) and Novo Nordisk Laureate (NNF19OC0056076) grants.

## METHODS

### Plant material and growth conditions

*Arabidopsis thaliana* ecotype Columbia (Col-0) was used as wild type (WT) control for all experiments. The mutant lines of *NKS1* (At4g30996), *nks1-1* (SALK_151073) and *nks1-2* (GK-228H05) were obtained from the Nottingham Arabidopsis Stock Centre (NASC) (Scholl et al., 2000) whereas *qua2-1*, *esmd1-1* were gifted by Gregor Mouille (INRAE, Paris) and Stephane Verger (UPSC, Umeå, Sweden), respectively. The various endomembrane compartment specific and other marker lines used in this study were obtained from NASC and/or obtained from original source are listed in Table S6.

Seeds were surface sterilized in 70% ethanol with 1% bleach for 5 minutes and washed with either water or 70% ethanol (5X) and sown on square petri plates of half concentration (2.2g/L) of Murashige and Skoog (MS) nutrient mix (Duchefa), 0.5% sucrose and 0.8% (w/v) plant agar (Duchefa) which was buffered to pH 5.8 by using 2.5mM 2-Morpholinoethanesulfonic acid (MES) (Sigma-Aldrich). The seeds were stratified at 4°C for 48 hours and grown in *in-vitro* growth chamber at 21°C under 16 hours of light and 8 hours of darkness (light grown seedlings).

For dark-grown experiments, after 48 hours of stratification, seeds were exposed to white light for 6 hours, then wrapped in aluminium foil and grown vertically in *in-vitro* growth chamber. This time point was used as start to count number of days after sowing (DAS). The growth period of seedlings varied in different experiments and details on this are mentioned in the respective figure legends.

After two weeks of growth in growth chambers, plants were transferred to 6 cm and/or 10 cm sized pots filled with peat substrate (made of peat, vermiculate and sand) with full nutrient supply (MPI *Arabidopsis* substrate, Stender, Germany) pre-watered with systemic fungicide (PREVICUR, Bayer Ltd, 1.5mL/L) and Boron solution (1mL/L). The plants were grown under long day conditions (16 hr light, 21°C, RH-50% and 8 hr dark, 17°C, RH-50%). Plants were genotyped using the primers indicated in Table S7 for *nks1* alleles and *qua2-1* T-DNA insertion lines. To genotype *esmd1-1*, we developed new dCAPS primers using (http://helix.wustl.edu/dcaps/; Neff et al., 2002) using the primers are listed Table S7. The resulting PCR fragment was subjected to restriction digestion by BseXI enzyme, generating two fragments of 500+302 bp in Col-0 and 800+1 bp in *esmd1-1*. The *esmd1-1* mutants were then confirmed by sequencing for the presence of single nucleotide polymorphism reported by Verger et al., 2016. To genotype *qua2-1*, we amplified a fragment using primers listed in Table S7 and allele was confirmed by sequencing for the presence of single nucleotide polymorphism as reported by Verger et al., 2018.

Various double mutants and marker lines used in this study was generated by crossing.

### Brightfield microscopy and histology

#### Hypocotyl length and time-lapse growth analysis

Three-day and six-day old dark grown seedlings were scanned using scanner (EPSON perfection V600 photo) at 800 dots per inch (dpi) resolution. Hypocotyl lengths were measured using a segmented line in Fiji (ImageJ) software. Time lapse hypocotyl growth kinetics was done according to Jonsson et al. (2021); briefly, seedlings were grown vertically on ½ MS media were imaged using Canon D50 camera at 1 hour interval without infrared filter, and hypocotyl growth was measured using Fiji.

#### Visualization of seed mucilage by Ruthenium Red staining assay

Seeds were incubated in Tris 10mM (pH 7.6) and shaken vigorously on an orbital shaker for 2 h at room temperature to hydrate and release mucilage from the epidermal seed coat. This solution was replaced with 800 μl of 0.01% ruthenium red solution (11103-72-3, Sigma-Aldrich). The seeds were again shaken vigorously on an orbital shaker for 1 h. The seeds were then washed with water to remove excess stain. The seeds suspended in water were then mounted in depression slide and imaged using a compound microscope (McFarlane et al., 2014).

#### Tissue integrity assay

Six-day-old etiolated seedlings were stained in aqueous solution of 0.05 % (w/v) Toluidine blue for 1 min. Then, seedlings were washed gently with water once and imaged immediately using a compound microscope (Tanaka et al., 2004; Neumetzler et al., 2012).

### *In silico* analyses

#### *NKS1* gene expression

*NKS1* (At4g30996) gene expression patterns were accessed via ePlant (https://bar.utoronto.ca/eplant/; Waese et al., 2017).

#### Coexpression analyses

The putative functional homologs of *NKS1* gene were identified using ATTED-II (https://atted.jp/; Obayashi et al., 2018) and are listed in Table S1.

#### Gene Ontology (GO) analyses

GO analyses were conducted via the Gene Ontology Resource interface (http://geneontology.org/) where the top 100 co-expressed genes were uploaded and compared against the Arabidopsis genome. The outputs are listed in Table S2.

#### Protein domain structure prediction

Predicted protein domain architecture was accessed via InterPro (https://www.ebi.ac.uk/interpro/about/interpro/; Blum et al., 2021). Transmembrane spanning helices were predicted using the TMHMM Server v.2.0 (http://www.cbs.dtu.dk/services/TMHMM/; Krogh et al., 2001).

#### Phylogenetic analyses

NKS1 amino acid sequence was used to search homologues against publicly available database such as PLAZA (Van Bel et al., 2017), NCBI (https://www.ncbi.nlm.nih.gov/), and Phytozome (Goodstein et al., 2012). The identified protein sequences from Arabidopsis thaliana (At4g30996, At2g24290, At4g04360 and At2g32580), *Oryza sativa* (Os04g42340,Os03g56610), *Brachypodium distachyon* (BdiBd21-3.2G0728900.1, BdiBd21-3.5G0190400.1,BdiBd21-3.1G0094300.1), *Populus trichocarpa* (Potri.006G187800,Potri.018G111100,Potri.002G227100,Potri.004G006500, Potri.013G026900, Potri.005G038000, Potri.011G009200, Potri.011G009300, Potri.T040000), *Picea abies* (PAB00059623), *Marchantia polymorpha* (Mapoly0042s0084, Mapoly0138s0016), *Selaginella moellendorfii* (SMO358G0620), *Utricularia gibba* (UGI.Scf00506.16776, UGI.Scf00037.4142, UGI.Scf00027.3259) were used to construct phylogenetic tree according to Zhang et al. (2015). Amino acid sequences were aligned by MUSCLE algorithm at MEGA and subsequently phylogenetic tree was constructed using maximum-likelihood Le and Gascuel (LG) model. Parameter were used as phylogeny test-bootstrap method, No. of bootstrap replications-1000, Substitution type-amino acid, model-LG, rates among sites-Gamma distributed, No. of discrete Gamma categories-5.

### Gene expression assay by qRT-PCR and RT-PCR

Total RNA was isolated from six-day old etiolated seedlings of Col-0, *nks1-1* and *nks1-2* using RNeasy Plant mini kit (74904, QIAGEN). cDNA was synthesized from 1 μg total RNA using iScript cDNA synthesis kit (1706691, Bio-Rad). Transcript levels were analysed from three biological replicates by real time quantitative PCR (qRT-PCR). Quantitative expression of *NKS1* was determined in wild type and *nks1* alleles by qRT-PCR using SYBR Green (Applied Biosystems) reaction mixture on an ABI PRISM 7900 HT sequence detection systems (Applied Biosystems) (PCR reaction: 50°C for 2 min, 95°C for 10 min, 40 cycles of 95°C for 15 s, and 60°C for 1 min. Amplicon dissociation curves, i.e. melting curves, were recorded after cycle 40 by heating from 60°C to 95°C with a ramp speed of 1.9°C min^−1^) or CFX96 Touch Real-Time PCR detection system, Bio-Rad (PCR reaction:95°C for 5 min, 95°C for 10 sec, 60 °C for 10 sec, 72°C for 15 sec. The melting curve was recorded after 39 cycle by heating from 65°C to 95°C with increment of 0.5°C.

The relative expression values were calculated by the 2^-ΔΔCq method using Reference Gene Index (RGI). *POLYUBIQUITIN10* (At4g05320), *ACTIN2* (At3g18780), *PROTEIN PHOSPHATASE* (At1g13320) and SAND family protein encoding gene *SAND* (At2g28390) were used to calculate the reference gene index (Czechowski et al., 2005). The primers used to amplify *NKS1* and *FADlox* are given in Table S7.

Semi-quantitative RT-PCR was performed by intron spanning primers for NKS1 (RT1FP and RT1RP; Table S7) and APT1 gene was used as internal normalization and cDNA loading control. The 1:10 diluted cDNA prepared from 1μg of RNA was used from three biological replicates of Col-0, *nks1-1* and *nks1-2*. PCR was performed with following conditions: 94°C for 10 sec, 58°C for 30 sec, 72°C for 1 min.

### Bimolecular Fluorescence Complementation (BiFC)

#### Cloning BiFC constructs

NKS1 (At4g30996), QUA1 (At3g25140), QUA2(At1g78240), Got1P homolog (At3g03180) (Zhang et al., 2016) and IAA1 (At4g14560) (Pandey et al., 2018) were PCR amplified from Arabidopsis Col-0 cDNA using forward primer with attB1 and reverse primer with attB2 site; all primers are listed in Table S7. The PCR fragment was cloned into pDONR207 using BP Clonase Enzyme II (Cat No. 11789020, Thermo Scientific) mix at 25°C. Entry clones were then sub-cloned into BiFC specific destination vectors (pDEST-gwVYCE and pDEST-gwVYNE) (Gehl et al., 2009) using Gateway LR Clonase II Mix kit (Cat. No. 17791100, Thermo Scientific). The clones confirmed by PCR and restriction digestions.

#### Preparation and transient expression in *Arabidopsis* root suspension culture protoplasts

Arabidopsis root suspension culture were grown in 25 mL of Murashige and Skoog (MS) media (4.33 g/L MS Salts (Duchefa), 2 ml/L B5 vitamin stock, 3% sucrose, 0.24 mg/L 2,4-D, 0.014 mg/L ketenin, dissolved in de-ionized water and set pH to 5.7) at 22°C for 4 days. Protoplasts were isolated in enzyme solution (1% Cellulase (Onozuka R-10), 0.2% Macerozyme (Serva) in B5+0.34 M Glucose Mannitol (GM) solution (4.4 g/L MS (Duchefa), 30.5 g/L Glucose (VWR Chemicals), 30.5 g/L Mannitol (Sigma), pH to 5.5 with KOH) with slight shaking for 3-4 h and afterwards centrifuged and washed with B5+0.34M GM at 192g for 5 minutes. The pellet was dissolved in B5+0.28 M sucrose. 5μg of each plasmid was mixed in 50 μL of protoplast suspension and 150 μL of 25% PEG 6000 solution, then incubated in dark for 20 minutes and the reaction was stopped by addition of 500 μL 0.275 M Ca(NO_3_)_2_ followed by centrifugation for 1 minutes at 123g. Supernatant was discarded and 500 μL of MS+0.34 M GM was added to the cells which were then incubated in dark for 16h.

#### Confocal imaging

Transfected protoplasts were mounted under a coverslip separated from the slide with double sided tape and viewed with x20 or x40 (water immersion) objectives of Zeiss LSM880 confocal scanning microscope. The YFP fluorescence was excited at 514nm and emission spectra was detected in spectral range of 500-600nm. Lambda wavelength mode of imaging used to confirm peak signal emission spectra.

### GO-PROMPTO assay

#### Cloning & plant transformation

Modified GO-PROMTO assay (Søgaard et al., 2012) with VENUS as the fluorescent marker (Lampugnani et al., 2016) was used to determine the topology of NKS1. The NKS1 CDS was amplified with SfoI-forward primer and KpnI-reverse primer (Table S7) and cloned into an SfoI- and KpnI-linearized pSUR. The constructs generated were verified through sequencing and transformed into the AGL1 strain of *A. tumefaciens* by electroporation with the helper plasmid pSOUP. Transient expression in *N. benthamiana* leaves and imaging was carried out as previously described in (Sanchez-Rodriguez et al., 2018).

### Generation of NKS1-GFP/RFP translational fusion constructs

The coding sequence of NKS1 with or without stop codon was amplified by PCR using gene specific primers (Table S7) from cDNA using Phusion High-Fidelity DNA polymerase (F530S, NEW ENGLAND BioLabs, Inc) or PrimeSTAR HS DNA polymerase (R010A, Takara Clonetech Ltd). The fragments were introduced into pENTR-D-TOPO vector by pENTR^™^ Directional TOPO Cloning Kit (K2400-20, Life Technologies). pENTR-D-TOPO-NKS1 with or without stop codon was subsequently cloned into pUBN-GFP/RFP-NKS1 (N-terminal fusion) and pUBC-NKS-GFP/RFP (C-terminal fusion) plant expression vectors (Grefen et al., 2010) by Gateway LR clonase mix (11791-019, Life Technologies). The resultant constructs were transformed into *Agrobacterium tumefaciens* GV3101, which was used to transform Arabidopsis Col-0, *nks1-1*, and *nsk1-2* via floral dip (Clough SJ and Bent AF., 1998).

### Live cell imaging

#### Low water potential treatment and imaging of cell-cell adhesion defect phenotype

Water potential of ½ MS growth media was changed as described in Verger et al., 2018. Briefly, seedlings were grown on ½ MS media with 0.8% or 2.5% agar and hypocotyl growth was measured on six-day old dark grown plates. To check the cell adhesion defect, five-day old etiolated hypocotyls were stained with 0.2 mg/mL propidium iodide for 15 min. The seedlings were washed in water before imaging. The 3^rd^ or 4^th^ cell of basal part of hypocotyl from hypocotyl-root junction was used to image cell-adhesion defect (Verger et al., 2018). The seedlings were imaged using Zeiss LSM 780 or 880 confocal laser scanning microscope (25X objective, N.A. 0.8) with excitation of 514 nm and emission was detected in the range of 600-650nm. The images were analysed using Fiji software. Initially, images were processed such as background subtraction was done using rolling ball radius of 20 pixels. The Z-stack projection was performed using sum slices. All the acquisition and processing steps were similar in all genotypes.

#### Spinning disk microscopy

All other live cell imaging was conducted using CSU-X1 Yokogawa spinning disc head fitted to a Nikon Ti-E inverted microscope, a CFI APO TIRF X100 N.A. 1.49 oil immersion objective, an evolve charge-coupled device camera (Photometric Technology) and a X1.2 lens between the spinning disc and camera. GFP was excited at 491 nm and mCherry at 561 nm using a multichannel dichroic and an ET525/50M or an ET595/50M band pass emission filter (Chroma Technology) for GFP/YFP and mCherry fluorophores, respectively. Alternatively, live cell imaging was conducted with an inverted Nikon Ti-E with an Andor Revolution CSU-W1 spinning disk, an Andor FRAPPA photobleaching unit, two Andor iXon Ultra 888 EM-CCD cameras, and 100x or 60x N.A. 1.49 Apo TIRF oil-immersion objectives. GFP was excited with a 488 nm laser and emission collected with a 525/50 nm band pass filter; YFP was excited with a 515 nm laser and emission collected with a 535/30 nm filter; mCherry and RFP were excited with a 561 nm laser and emission collected with a 610/40 nm filter.

#### Sample preparation

For seedling imaging, 3-day-old etiolated hypocotyls or roots were mounted in water under a pad of 0.8% agarose (Bioline). To limit the time that seedlings spent mounted, no more than three cells per seedling were imaged in any experiment. For BFA-treatments, seedlings treated with BFA (50μM) diluted in ½ MS media with 1% sucrose for timepoints indicated and washout was performed with ½ MS media with 1% sucrose. FM4-64 straining was performed for 10 minutes with 2μM FM4-64.

#### Simultaneous dual wavelength imaging

For double Golgi marker colocalization and Golgi-TGN colocalization, both channels were excited simultaneously and emission was collected simultaneously using the excitation and emission parameters described above and two Andor iXon Ultra 888 EM-CCD cameras to eliminate time-lag between collecting two channels (collected at ~400ms exposure each), and potential displacement due to the rapid cytoplasmic streaming of Golgi bodies and TGN (up to 4.2 μm/s; Nebenführ et al., 1999). z-stacks were collected with 0.2 μm spacing using the 100x N.A. 1.49 objective.

### Live cell image analyses

All image processing was performed using Fiji software. For analysis involving measurement of signal intensity, only linear adjustment were made. For other images, background signal was reduced using the ‘Subtract Background’ tool (rolling ball radius of 20 to 30 pixels). Image drift was corrected using the Fiji plugin StackReg (Thevenaz et al., 1998).

#### Co-localization

Co-localization between NKS1-GFP or NKS1-RFP and compartment marker lines was analysed as described by (Gendre et al., 2011; Boutte et al., 2013; Gendre et al., 2013). All images were analysed using JACoP plugin in Fiji (Bolte and Cordelieres, 2006) using the appropriate hardware settings from the microscope, but otherwise default parameters.

#### Simultaneous dual wavelength imaging

Images from the two cameras were aligned relative to a calibration slide, then regions of interest were selected from 3 cells per seedling for colocalization analysis. Colocalization was quantified from z-stacks (with 0.2 μm spacing using the 100x N.A. 1.49 objective) using the DiAna plugin for Fiji (Gilles et al., 2017) with the following parameters for DiAna-Segment: no filtering, manual thresholding, object size >10 pixels and then DiAna-Analyze was used to measure centre-centre distance between the segmented objects.

#### CESA speed and density measurements

Wild type and *nks1-2* GFP-CESA3 seedlings were imaged with 10 sec time intervals for 600 sec. Background signal was reduced using the ‘Subtract Background’ tool (rolling ball radius of 20 to 30 pixels). If necessary, image drift was corrected using the Fiji plugin: StackReg (Thevenaz et al., 1998). CESA speed and density were determined according to Sampathkumar et al. (2013).

#### Golgi speeds

Golgi movement was tracked using Fiji-TrackMate (Tinevez et al., 2017). Golgi were detected as particles of 10 pixels and then linked in different frames using simple linear assignment problem tracker with a maximum linkage distance of 15 pixels, a maximum gap closing distance of 15 pixels, and a maximum frame gap number of 3. The parameter “Mean Speed” was used to calculate the average Golgi motility rate.

#### FM4-64 internalization

Wild type and *nks1-2* roots were treated with BFA (50μM) for 30 minutes and FM4-64 (2μM) for 10 minutes, then mounted and imaged. Maximum fluorescence intensities of BFA bodies were measured and compared to the plasma membrane intensities using Fiji (Gadeyne et al., 2014).

#### PIN2-GFP recycling

For PIN2-GFP quantification, plasma membrane signal was measured using a segmented line drawn along the apical surface of the plasma membrane of a cell and intracellular signal was measured within a hand-drawn polygon and the mean signal intensities were measured in Fiji and used to calculate the ratio of plasma membrane:intracellular signal. The number of BFA bodies within this polygon were manually counted then its area was measured in Fiji and used to calculate the number of BFA bodies per unit area.

### Cell wall analyses

Cell wall analyses were conducted on 6-day-old etiolated seedlings of Col-0, *nks1-1*, *nks1-2*. The seedlings were harvested in 70% ethanol and the seed coats were removed carefully from seedlings. The cleaned seedlings were collected in 2 mL Eppendorf tube and washed with 70 % ethanol. The ethanol was completely evaporated by drying seedlings at 60° C overnight. The dried samples were grounded by Retsch Mill (Retsch Inc.) for 3 minutes at 30 Hz, then washed once with 70% ethanol and ground for 1 more min. The content was vortexed thoroughly and spun down at 14,000 rpm for 10 min. The pellet was then washed twice with chloroform: methanol (1:1 v/v). The resultant pellet was washed with acetone, and dried overnight to obtain cell wall material (CWM).

#### Preparation of samples via acid hydrolysis

0.5 to 1 mg of dried CWM was weighed in screw capped Eppendorf tubes. An internal standard (30μg of inositol) was added to the CWM. 250 μL of 2M trifluoroacetic acid (TFA) was added to the pellet and incubated at 121°C for 1 hour in a heating block. The TFA was later evaporated by washing with isopropanol thrice under a steam of dried air. 300 μL of water was added to the pellet and mixed by vortexing followed by sonication. The mix was spun down at 14,000 rpm for 10 min. The pellet was dried overnight and used by cellulose estimation for modified Seaman analysis (Selvendran and O’Neill, 1987).

#### Estimation of cellulose content

The hexose content was estimated by the Anthrone assay (Updegraff, 1969). The pellet was dissolved in 175 μL of 72% sulphuric acid by shaking and vortexing. Then 425 μL of water was added and mixed well. 100 μL of sample was used to estimate hexose content. 200 μL of 0.2% anthrone reagent was added to the sample. Glucose standards were prepared along with the sample. The content was boiled at 121°C for 5 min. 200 μL of sample was loaded into a microtiter plate and absorbance was measured at 640 nm in photometer. The cellulose content in sample was calculated from D-Glu standard curve.

#### Monosaccharide analysis

Alcohol insoluble residue (AIR) was prepared, and pectin and hemicellulose fractions were extracted sequentially as previously described (Rautengarten et al., 2019). AIR (10-15 mg) obtained from dark-grown hypocotyls was extracted using 50 mM CDTA (pH 6.5), 50 mM Na_2_CO_3_, 1 M and 4 M KOH for 2 h at room temperature and 16 h at 4°C. CDTA fractions were combined and dialyzed against 50 mM sodium acetate, pH 5.5. Sodium carbonate fractions were combined and neutralized with acetic acid. The 1 M KOH and 4 M KOH fractions were similarly combined and neutralized with concentrated HCL. All the fractions were further dialyzed three times 16 h at 4°C against de-ionized water and lyophilized. The remaining insoluble residue (pellet) was washed with water, 70% (v/v) ethanol, and acetone and dried. Total alcohol insoluble residue or corresponding cell wall fractions were hydrolyzed in 2 N trifluoroacetic acid (TFA) for 1 h at 120°C. High-Performance anion exchange chromatography (HPAEC) coupled with pulsed amperometric detection (PAD) was performed on an ICS 5000 device (Dionex) using a CarboPac PA20 (3 × 150 mm) anion-exchange column (Dionex) (Rautengarten et al., 2016).

### Immunoprecipitation of NKS1-GFP and interactors

#### Protein extraction

7-day-old NKS1-GFP seedlings were grown in liquid ½ MS pH 5.8 with 1% sucrose and snap frozen in liquid nitrogen, freeze-dried for 5 hours in a LSCplus Freeze Drier (Christ), and stored at −80°C until further analysis. 3 independent experiments of 3 (E1, E2) or 2 (E3) biological replicates (1 flask of seedlings for each replicate) were performed for a total of 8 biological replicates. Dried samples were ground with mortar and pestle in liquid nitrogen. Total protein was extracted using an extraction buffer comprised of 50 mM MOPS (M1254, Sigma) pH 7.0, 2 mM EDTA (798681-1KG, Sigma), 2 mM EGTA (E4378-25G, Sigma), along with 1 tablet of cOmplete^™^ EDTA-free protease inhibitor cocktail (5056489001, Roche) per 50 mL of buffer. The homogenized solution was centrifuged at 3,270 g in an Optima L-80 Ultracentrifuge with a 70.1 Ti rotor (Beckmann). The supernatant was centrifuged again for 1 h at 100,000g in the same equipment to extract the microsomes. The microsomal pellet was resuspended in pellet buffer of 10 mM Tris-HCl (Sigma) pH 7.5, 150 mM NaCl, 0.5 mM EDTA and 0.5% NP-40. Protein was quantified using Pierce Bicinchoninic Acid (BCA) assay kit (Thermo). The samples were adjusted to get at least 480 μg of protein. The next steps were performed on ice. GFP-Trap A beads (CT-gta-20, BioNovus Life Sciences) were conditioned with two washes using a dilution buffer that consisted of 10 mM Tris-HCl pH 7.5, 150 mM NaCl, 0.5 mM EDTA. Proteins were bound to the beads by adding 500 μL of microsomal fraction proteins in pellet buffer and samples were tumbled for 1h at 4° C. Then, they were centrifuged at 2,500 g for 2 minutes at 4°C. Samples were washed with 500 μL of pellet buffer three times, and twice with dilution buffer. Then, 25 μL Elution buffer I, consisting of 50 mM Tris-HCl pH 7.5, 2 M urea, 5 μg/mL Mass Spec Trypsin/Lys-C Mix (V5073, Promega) and 1 mM DTT was added to the samples, and incubated in a thermomixer at 30 °C at 400 rpm for 30 min. Then, samples were centrifuged at 2,500x g for 2 minutes at 4°C, and the supernatants were transferred to a fresh vial. A second and third elution was made using 50 μL of 50 mM Tris-HCl pH 7.5, 2 M urea, 5 mM iodoacetamide (I6125, Sigma) buffer, twice. The samples were dried on RVC 2-33 CDplus (John Morris) speed-vacuum evaporator for 1.5 h. The dried samples were stored at −20°C.

#### Mass spectrometry

Samples were analyzed using liquid chromatography with tandem mass spectrometry (LC MS/MS) on Orbitrap Fusion Lumos Tribrid mass spectrometer and an Ultimate 3000 RSLC nano system (Thermo Scientific). Dried samples were resuspended in 5% acetonitrile, 1% trifluoroacetic acid; 6 μL of sample were injected for pulldown fractions. The samples were trapped on a PepMap 300 μm × 1 mm (C18, 5 μm, 100 Å) precolumn and separated on an Acclaim PepMap RSLC 75 μm × 50 cm (C18, 2 μm, 100 Å) column (Thermo Scientific). The flow rate was 220 nL·min^−1^ with a total analysis time of 90 minutes. The experiments were performed using a nano electrospray ionization source at positive mode and Fusion Lumos Orbitrap mass spectrometer (Thermo Fisher Scientific). The mass spectrometry data was acquired on for one full scan MS mode and as many data dependent HCD-MS/MS spectra as possible. All mass spectrometry data were acquired using Orbitrap mass analyzer.

#### Data analysis

Peptide identification was performed on Proteome Discoverer (v.2.3 Thermo) searching the Arabidopsis database (TAIR10) and label free quantification was performed using the Minora Feature Finder. A total of 2,760 proteins were identified. Statistical analysis was performed using RStudio (v3.6.2). Interaction candidates were quality filtered based on the following criteria: ≥10% coverage, ≥ 2 peptides, FDR ≤ 0.05, Score ≥ 10, ≥ 1 unique peptide, resulting in 407 proteins. To select the candidates the samples were further filtered selecting for those with an average abundance across all experiments ≥100, found with ‘high’ confidence in all rounds, and present in at least 6 samples, resulting in 248 proteins. We then used SUBA (Hooper et al., 2017) to narrow down potential candidates. AGIs corresponding to the proteins identified above the quality control thresholds described were input to SUBA and filtered for “Location All Predictors” containing the term “golgi”. This generated a very loose filter for any proteins that were identified as possibly Golgi-localized by any predictor curated by SUBA, resulting in 94 proteins. To identify high confidence candidates, the resultant list was cross-referenced with 3 others, published Golgi proteomes (Nikolovski et al., 2012; Parsons et al., 2012; Parsons et al., 2019). The final candidates list (Table S4) includes 37 proteins that appear at least once in one of the Golgi proteomes, while also detected in the NKS1 pulldown in all three experiments and at least six out of eight biological replicates.

### Transmission electron microscopy (TEM) & transmission electron tomography (ET)

For both TEM and tomography, etiolated 3-day-old seedlings were cryofixed using a Leica HPM-100 high pressure freezer using 1-hexadecene as a cryoprotectant and B-type carriers, according to McFarlane et al. (2008). Freeze-substitution was performed in in 2% (w/v) osmium tetroxide (Electron Microscopy Sciences) and 8% 2,2-dimethoxypropane (w/v) in anhydrous acetone using a Leica AFS2 automatic freeze substitution unit at −85°C for 4 days, then the temperature was gradually raised to room temperature over 2 days. Samples were washed 5 times with anhydrous acetone, then infiltrated with Spurr’s Resin (Electron Microscopy Sciences) over the course of 4 days. Resin-infiltrated samples were polymerized at 65°C for 36 hours.

#### TEM

~80 nm thick (silver) sections were cut using a UC7 Ultramicrotome and a DiATOME diamond knife, placed on Gilder fine bar hexagonal 200 mesh grids coated with 0.3% formvar (Electron Microscopy Sciences). Grids were post-stained with 1% aqueous uranyl acetate (Polysciences) and Sato’s triple lead (sodium citrate, lead acetate, lead citrate from BDH, lead nitrate from Fisher) and imaged with a Phillips CM120 BioTWIN transmission electron microscope with a Gatan MultiScan 791 CCD camera and a tungsten filament at an accelerating voltage of 120 kV. Golgi features of genotype-blinded images were manually measured in Fiji.

#### ET

5 serial sections ~250-300 nm thick (green) were cut were cut using a UC7 Ultramicrotome and a DiATOME diamond knife, placed on Maxtaform copper/rhodium 2×1 mm slot grids coated with 0.8% formvar (Electron Microscopy Sciences), post-stained 1% aqueous uranyl acetate (Polysciences) and Sato’s triple lead (sodium citrate, lead acetate, lead citrate from BDH, lead nitrate from Fisher), and then coated with 15 nm colloidal gold (Ted Pella) as a fiducial marker. Samples were imaged with a FEI Tecnai F30 transmission electron microscope at an accelerating voltage of 300 kV equipped with a CETA 4×4k CMOS camera. Dual-axis tomograms were collected in a tilt range of +65° to −65° with 2° tilt steps per image per image thereafter using FEI Explore3D automated tomography acquisition package with manual rotation of the sample by ~90° between collection of the two tilt axes. Tomograms were aligned, reconstructed, and modelled in Etomo and IMOD (Kremer et al., 1996). In Etomo, fiducial gold on both sides of the section were used for alignment and 22 iterations of SIRT (Simultaneous Iterative Reconstructive Technique) was used for reconstruction. Dual-axis tomograms were aligned using fiducial markers from both tomograms then combined using Etomo. Tomograms were manually segmented using IMOD; Golgi cisternae and TGN were modelled manually as closed objects every 3 slices without interpolation, while vesicles were modelled as spherical scattered objects seeded at the section in which they displayed maximum diameter; after modelling, all surfaces were meshed and equally smoothed using the smoothsurf function in IMOD over 12 sections.

### Cryo scanning electron microscopy

Etiolated 3-day-old seedlings were processed according to McFarlane et al. (2021). Seedlings were mounted in Tissue-Tek (Sakura-Finetek) on a sample holder, plunge-frozen in a liquid nitrogen slush (roughly −210°C), then transferred to a Gatan cryostage. Ice crystals were evaporated at −95°C for 2.5 minutes, coated with 60:40 gold-palladium alloy for 120 sec (about 6 nm) under argon at −120°C, and then transferred into the FEI Quanta cryo scanning electron microscope. To ensure that cell separation was not an artefact of sample preparation, wild type, *nks1-1*, and *nks1-2* seedlings were mounted on the same sample holder and processed together. Stage temperatures were maintained below −120°C while images were collected at accelerating voltage of 5 kV and working distance of 10 mm using the E-T detector.

### Statistical analyses

Statistical tests were conducted using SPSS (IBM Corp.); sample sizes and details of each statistical test are presented in figures and figure legends.

## SUPPLEMENTAL MATERIAL

**Figure S1: *NKS1* gene structure and characterization of *nks1* alleles.**

(A) Schematic map of the *NKS1* (At4g30996) gene, with position of *nks1-1* and *nks1-2* allele T-DNA insertions indicated. Black and white box indicate coding and non-coding regions, respectively and lines indicate introns. Green arrow shows position of primers used to amplify NKS1 gene from CDNA. qRT2FP and qRTFP2RP (purple arrows) were used to perform quantitative real time PCR and whereas RT1FP and RT1RP (red arrows) were used to perform semi-quantitative RT-PCR. (B) Semi-quantitative RT-PCR analysis of *NKS1* transcript levels in three biological replicates of Col-0, *nks1-1* and *nks1-2* with *APT1* as a control. All primers used for this study are listed in Table S7. (C) ePlant eFP viewer (Waese et al., 2017) representing *NKS1* expression throughout the different Arabidopsis tissues depicted, according to the indicated colour scale.

**Figure S2: Functional NKS1-GFP is localized to Golgi apparatus and not the TGN or other endomembrane compartments.**

(A) Representative confocal images showing co-localization of NKS1-GFP/RFP with markers for the endoplasmic reticulum (GFP-HDEL), *trans*-Golgi network (TGN; VHAa1-mRFP), and late endosomes (RFP-WAVE7 and RFP-WAVE2) in 3-day old etiolated hypocotyl epidermal cells. (B) Representative images of NKS1-GFP co-localization with Golgi marker (XYLT-mRFP) or TGN marker (VHAa1-mRFP) after 60-minute BFA-treatment of 3-day old root epidermal cells. Scale bars represent 5μm in (A) and (B).

**Figure S3: NKS1 is a plant specific, transmembrane DUF1068 protein.**

(A) Phylogenetic analysis of full length NKS1 protein family. Amino acid sequences from *Arabidopsis thaliana* (4), *Populus trichocarpa* (9), *Oryza sativa* (2), *Brachypodium distachyon* (3), *Picea abies* (1), *Marchantia polymorpha* (2), *Selaginella moellendorfii* (1), and *Utricularia gibba* (1) were used for sequence alignment and tree construction. The clustered associated taxa are shown in percentage near the branches. The tree is constructed to scale with branch length measured in number of substitutions per site. (B) NKS1 is type II transmembrane protein as predicted by TMHMM Server v. 2.2; CBS Denmark. (C) Representative images of *N. benthamiana* leaf epidermal cells transiently transformed with V(I152L)N-ST, lumen facing reporter. Green signal in VC-NKS1+V(I152L)N-ST indicates that two proteins are on the same side of Golgi membrane. (D) Schematic representation of various domains known in NKS1 protein sequence predicted via InterPro (https://www.ebi.ac.uk/interpro/). NKS1 amino acid directs to IPR010471 entry at InterPro. Numbers represents positions of various domains in NKS1 protein sequence, TM-transmembrane domain (PHOBIUS, TMHMM entry); SignalP-TM (SIGNALP-GRAM_POSITIVE entry); CC-Coil (COILS entry), cytoplasmic and non-cytoplasmic domains (PHOBIUS entry).

**Figure S4: Golgi function is impaired in *nks1* mutants, but TGN structure and function is unaffected by loss of NKS1.**

(A) Representative confocal images of ratiometric SecGFP Col-0 and *nks1-2* hypocotyl epidermal cells of 3-day-old etiolated seedlings. (B) Representative images of simultaneous dual wavelength localization of Golgi (WAVE18) and *trans*-Golgi Network (VHAa1) dual markers in Col-0 and *nks1-2* hypocotyl epidermal cells of 3-day-old etiolated seedlings and linescan graph showing distance between dual markers in Col-0 and *nks1-2* from single Golgi-TGN object. (C) Representative transmission electron microscopy images of Golgi and TGN ultrastructure from Col-0, *nks1-1* and *nks1-2* hypocotyl epidermal cells of 3-day-old etiolated seedlings. (D) Representative images of PIN2-GFP localization in Col-0 and *nks1-2* root epidermal cells 3-day-old etiolated seedlings before and after BFA treatment, and following washout for 2h and 3h and quantification of plasma membrane:intracellular signal and number of BFA bodies as measurements of PIN2-GFP re-secretion from the TGN/BFA body. (E) Representative confocal images of Col-0 and *nks1-2* root epidermal cells of 3-day-old seedlings stained with FM-4-64 after BFA treatment and quantification of fluorescence intensities of FM-64 labelled bodies normalized to the fluorescence intensity of FM4-64 present in the plasma membrane. Statistically significant differences were determined by unequal variance, two tailed Student’s t test, where ns indicates non-significance. In violin plots, data distribution is outlined by the shape, plot box limits indicate 25^th^ and 75^th^ percentiles, whiskers extend to 1.5 times the interquartile range, median is indicated by a horizontal line, mean by a red dot and individual data points are shown, and n (cells, no more than 3 cells imaged per seedling) is indicated in parentheses. Scale bars represent 10 μm in (A), (B), (D) and (E) and 200 nm in (C)

**Figure S5: *nks1* mutants are defective in cell wall pectins, but cellulose synthesis is not significantly affected in *nks1* mutants.**

(A) Representative images of toluidine blue-stained Col-0, *nks1-1* and *nks1-2* 6-day-old etiolated seedlings. (B) Measurement of cellulose content from Col-0, *nks1-1* and *nks1-2* 6-day-old etiolated seedlings. (C) Representative images and quantification of GFP-CESA3 in Col-0 and *nks1-2* 3-day-old etiolated hypocotyl epidermal cells. Data are represented as single frames from the time lapse, a sum projection of the data (from 10-minute time lapse with 10 second intervals) and a kymograph from the yellow line indicated in the single frame. CESA speeds in the plasma membrane were quantified using the kymographs while CESA density at the plasma membrane was quantified from single frames. Statistically significant differences were determined by unequal variance, two tailed Student’s t-test, where ns indicates non-significance. In violin plots, data distribution is outlined by the shape, plot box limits indicate 25^th^ and 75^th^ percentiles, whiskers extend to 1.5 times the interquartile range, median is indicated by a horizontal line, mean by a red dot and individual data points are shown, and n is indicated in parentheses. Scale bars represent 500 μm in (A) and 10 μm in (C).

**Figure S6: *nks1* mutants phenocopy pectin synthesis mutants, *qua1*, *qua2*.**

(A) Quantitative reverse-transcription polymerase chain reaction (qRT-PCR) of *FADLox* transcript levels normalized to *UBQ10* from Col-0, *nks1-1* and *nks1-2*; bars represent means of three biological replicates ±SD. (B) Representative images of Col-0 and *nks1-2* seedlings grown on ½ MS growth medium supplemented with 0.5% sucrose (control) and 5% sucrose. (C) Representative z-projections (sum averages) of confocal stacks from propidium iodide-stained etiolated five-day old hypocotyl epidermal cell files from Col-0 and *nks1-2* grown on ½ MS growth medium supplemented with 0.8% agar (control) and 2.5% agar, and quantification of hypocotyl lengths of six-day old etiolated seedlings from Col-0, *nks1-1* and *nks1-2* grown on ½ MS growth medium supplemented with 0.8% agar (control) and 2.5% agar. Letters in (A) specify statistically significant differences among samples as determined by one way ANOVA followed by Tukey’s HSD test (p < 0.05). Statistically significant differences were determined by unequal variance, two tailed Student’s t-test, where ns indicates non-significance. In violin plots, data distribution is outlined by the shape, plot box limits indicate 25^th^ and 75^th^ percentiles, whiskers extend to 1.5 times the interquartile range, median is indicated by a horizontal line, mean by a red dot and individual data points are shown, and n is indicated in parentheses. Scale bars represent 500 μm in (B) and 200 μm in (C).

**Table S1: Genes co-expressed with NKS1 (At4g30996).** Data are from ATTED-II using At4g30996 as query gene.

**Table S2: GO enrichment analysis using the top 100 co-expressed genes from Table S1.** Output explanation may be found at: http://geneontology.org/

**Table S3: Monosaccharide composition of sequentially extracted and total cell wall material from wild type and *nks1-1 and nks1-2* mutants.** Values are shown as mole percent (mol%) of evaluated sugars. The control values are mean (s.d.) of three biological replicates (*n*=3). The *p*-values between the control and average were calculated using a Student’s t-test and significant differences (*p*<0.05) are marked (*).

**Table S4: Proteins identified in NKS1-GFP immunoprecipitation experiments.**

**Table S5: Additional Golgi and TGN measurements from TEM.** Golgi and TGN from morphometrics as quantified from transmission electron microscopy of high-pressure frozen, freeze-substituted hypocotyl epidermal cells of 3-day-old etiolated seedlings of Col-0 and *nks1-1* and *nks1-2*. Bold indicates statistically significant differences (p>0.05, t-test for measurements; p>0.01, χ^2^ test for proportions) between wild type and *nks1* mutants.

**Table S6: A list of all Arabidopsis seed lines employed in this study.** uNASC is The European *Arabidopsis* Stock Centre, Nottingham; GABI-KAT is from University of Bielefeld Germany.

**Table S7: A list of all synthetic oligonucleotides used in this study.** Oligonucleotides used in PCR-based experiments were ordered from MWG-Biotech AG (Germany) or Sigma (Australia).

