## Supplementary figures and images for "A DUF1068 protein acts as a pectin biosynthesis scaffold and maintains Golgi morphology and cell adhesion in Arabidopsis"

### Supplemental Figure 1

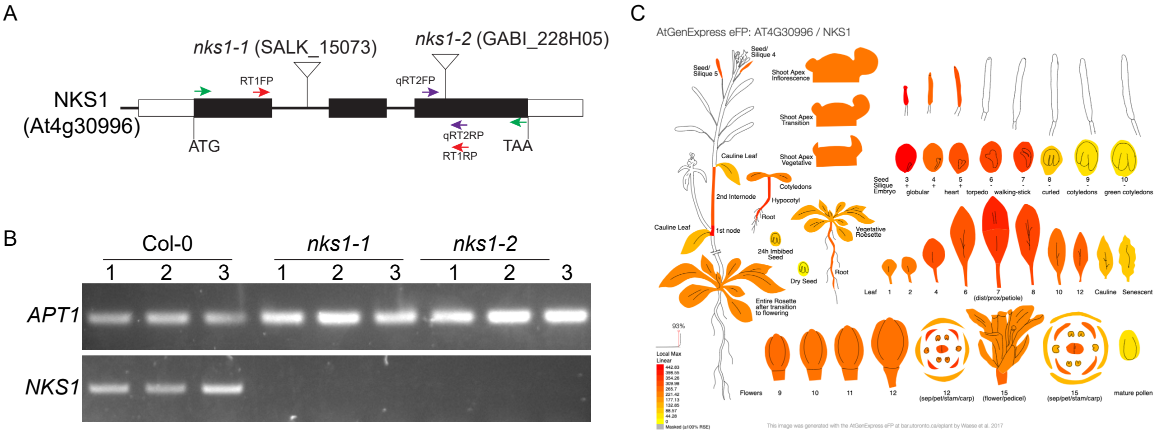


Figure S1: *NKS1* gene structure and characterization of *nks1* alleles.

### Supplemental Figure 3

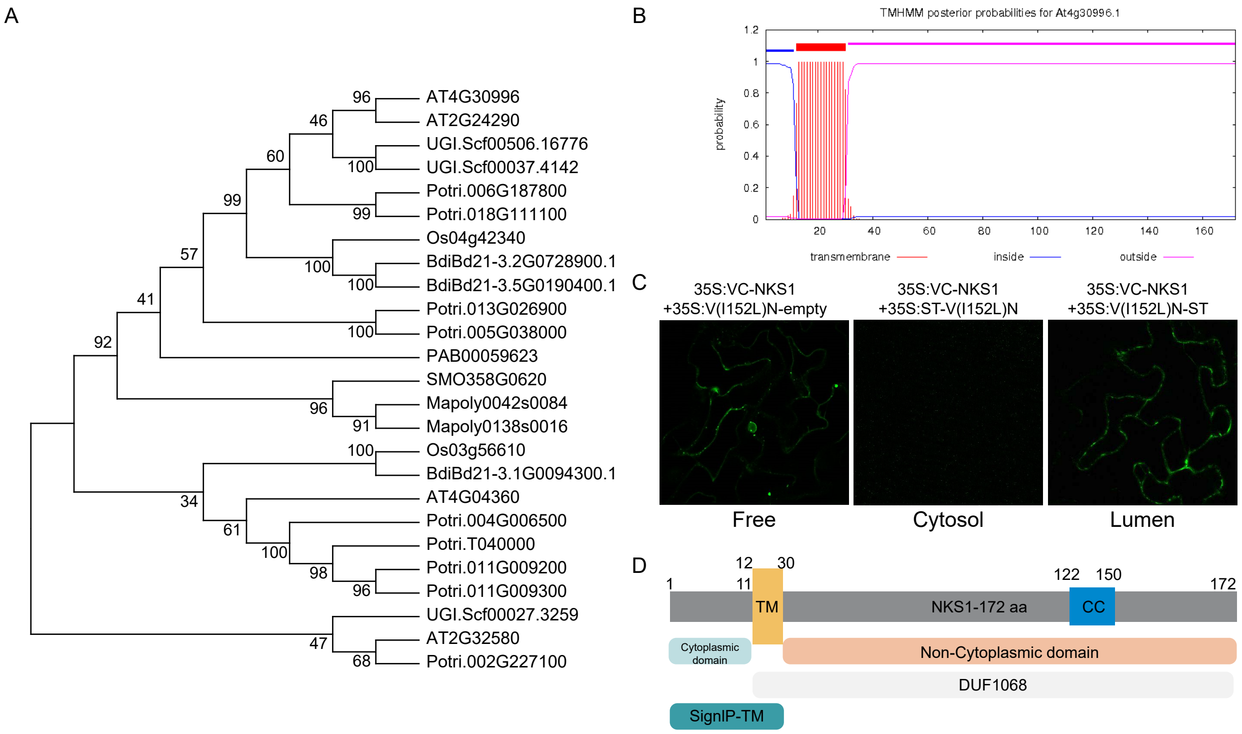


Figure S3: NKS1 is a plant specific, transmembrane DUF1068 protein.

### Supplemental Figure 6

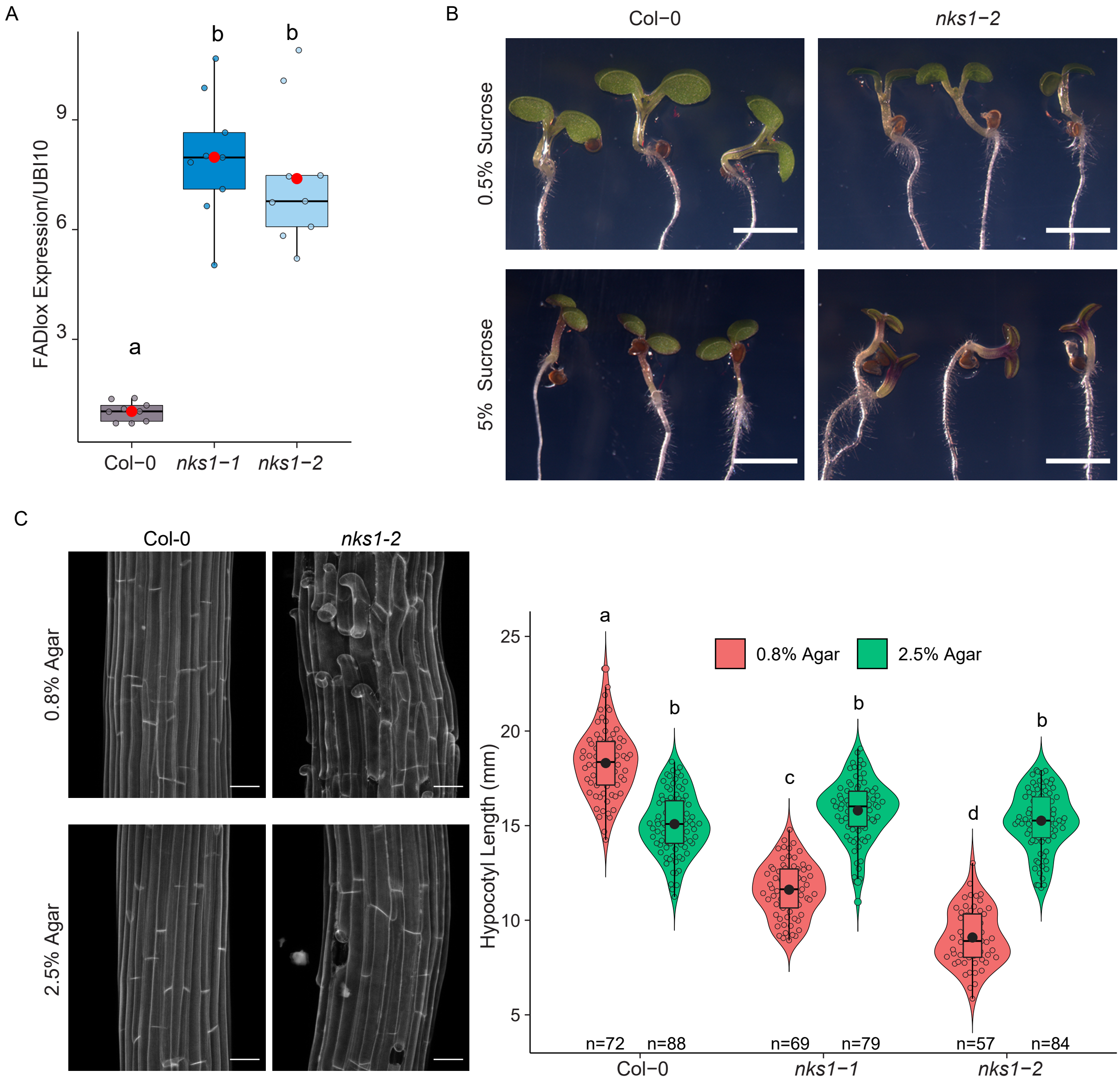


Figure S6: *nks1* mutants phenocopy pectin synthesis mutants, *qua1*, *qua2*.
