## Supplemental Figure 2 for "A DUF1068 protein acts as a pectin biosynthesis scaffold and maintains Golgi morphology and cell adhesion in Arabidopsis"

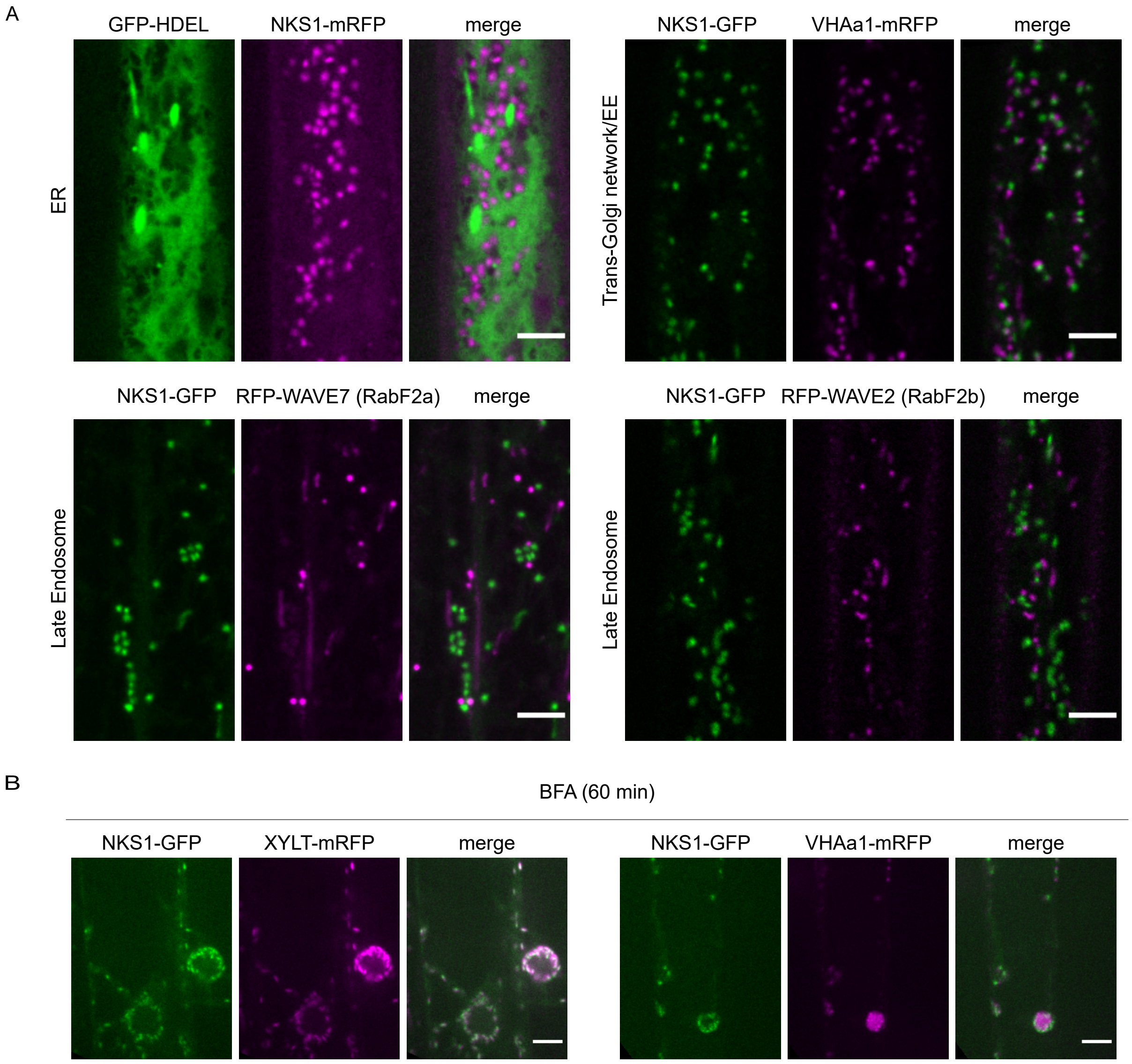


Figure S2: Functional NKS1-GFP is localized to Golgi apparatus and not the TGN or other endomembrane compartments.
