## Supplemental Figure 4 for "A DUF1068 protein acts as a pectin biosynthesis scaffold and maintains Golgi morphology and cell adhesion in Arabidopsis"

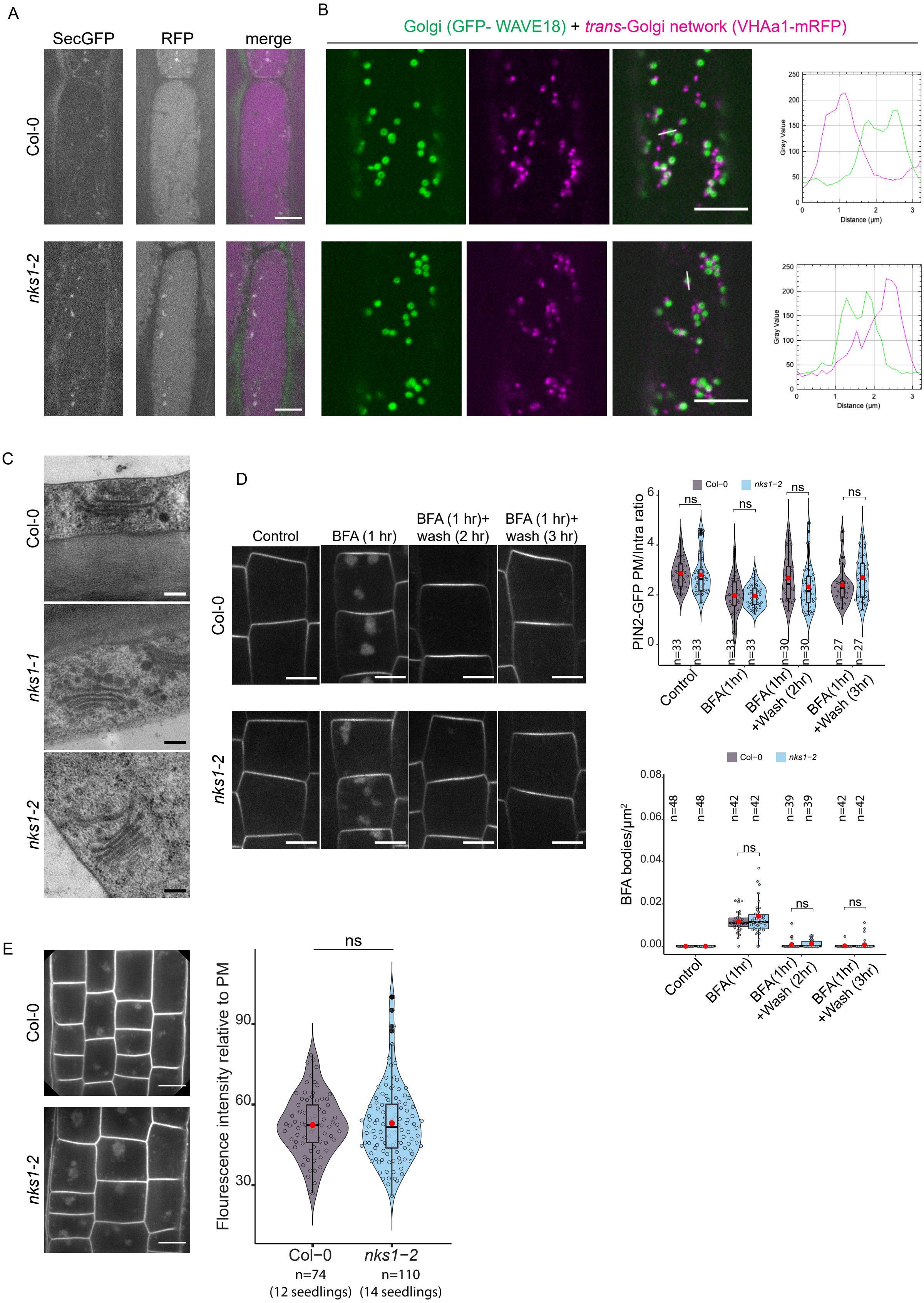


Figure S4: Golgi function is impaired in *nks1* mutants, but TGN structure and function is unaffected by loss of NKS1.
