## Supplemental Figure 5 for "A DUF1068 protein acts as a pectin biosynthesis scaffold and maintains Golgi morphology and cell adhesion in Arabidopsis"

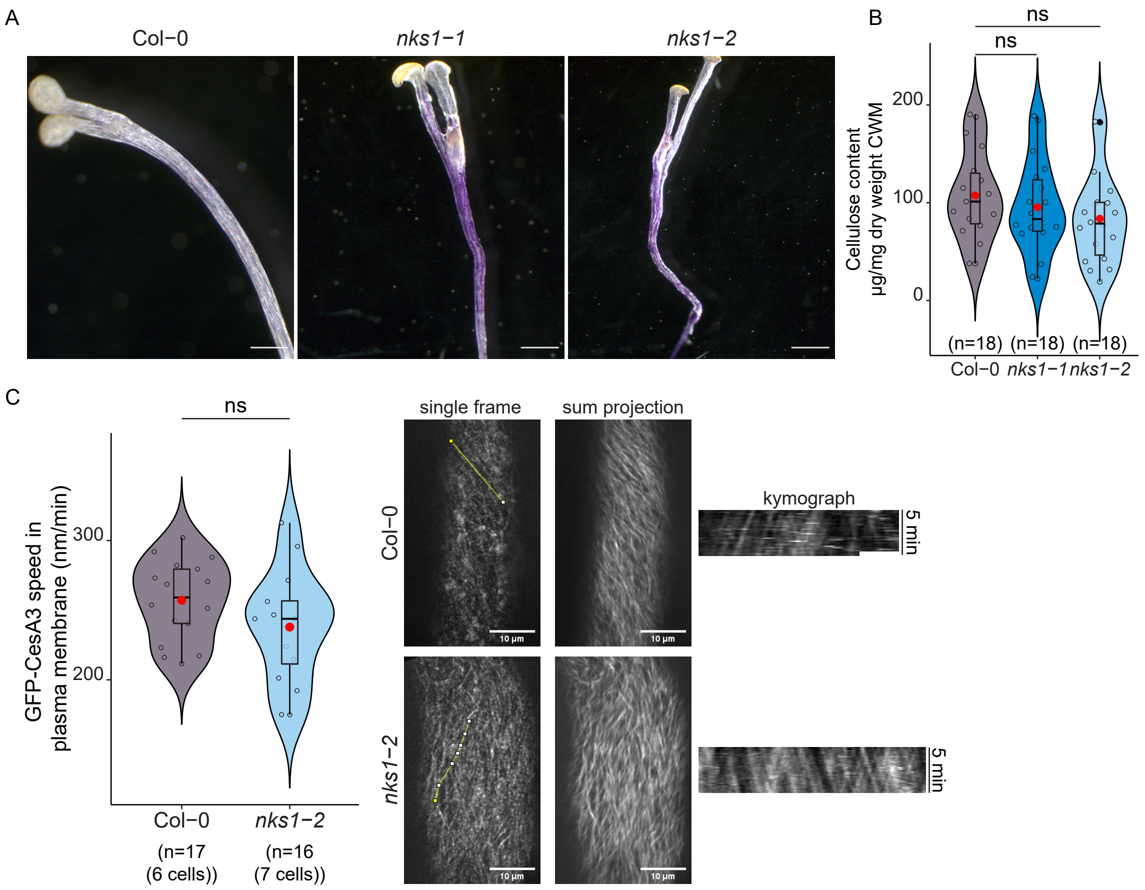


Figure S5: *nks1* mutants are defective in cell wall pectins, but cellulose synthesis is not significantly affected in *nks1* mutants.
