## Supplemental Table 3 for "A DUF1068 protein acts as a pectin biosynthesis scaffold and maintains Golgi morphology and cell adhesion in Arabidopsis"

**Table S3: Monosaccharide composition of sequentially extracted and total cell wall material from wild type and *nks1-1 and nks1-2* mutants.** Values are shown as mole percent (mol%) of evaluated sugars. The control values are mean (s.d.) of three biological replicates (*n*=3). The *p*-values between the control and average were calculated using a Student's t-test and significant differences (*p*<0.05) are marked (*).

| **Fraction** | **Fuc**  Avg mol% | **Rha**  Avg mol% | **Ara**  Avg mol% | **Gal**  Avg mol% | **Glc**  Avg mol% | **Xyl/Man**  Avg mol% | **GalA**  Avg mol% | **GlcA**  Avg mol% |
| --- | --- | --- | --- | --- | --- | --- | --- | --- |
| **CDTA**  wild type  *nks 1-1*  *p-*value  *nks 1-2*  *p-*value | 3.00 (0.18)  3.96 (0.13)*  0.002  3.99 (0.38)*  0.02 | 6.58 (0.22)  6.93 (0.10)  0.07  7.54 (4.09)  0.70 | 23.67 (0.28)  23.25 (0.25)  0.12  23.52 (2.53)  0.92 | 33.54 (1.80)  31.54 (0.84)  0.16  29.61 (1.82)  0.06 | 6.41 (2.12)  9.08 (5.21)  0.19  10.96 (0.10)*  0.02 | 5.11 (0.30)  5.21 (0.19)  0.65  5.25 (0.23)  0.57 | 21.68 (0.32)  19.52 (0.80)*  0.01  18.47 (0.40)*  0.0004 | 0.70 (0.05)  0.76 (0.00)  0.24  0.99 (0.39)  0.40 |
| **Na_2_CO_3_**  wild type  *nks 1-1*  *p-*value  *nks 1-2*  *p-*value | 1.63 (0.11)  1.47 (0.02)  0.06  1.47 (0.12)  0.15 | 7.75 (0.38)  7.52 (0.19)  0.40  8.99 (1.74)  0.29 | 14.84 (0.70)  16.58 (0.40)*  0.02  16.46 (0.53)*  0.03 | 56.56 (1.10)  57.33 (0.39)  0.32  53.74 (3.79)  0.28 | 1.31 (0.38)  1.43 (0.15)  0.66  3.35 (1.68)  0.11 | 3.18 (0.40)  3.01 (0.29)  0.57  3.09 (0.82)  0.86 | 14.53 (1.09)  12.62 (0.60)  0.06  12.57 (0.89)  0.07 | 0.18 (0.24)  0.05 (0.03)  0.38  0.33 (0.48)  0.66 |
| **1 N KOH**  wild type  *nks 1-1*  *p-*value  *nks 1-2*  *p-*value | 3.60 (1.24)  3.53 (1.77)  0.44  3.79 (0.62)  0.82 | 2.73 (0.16)  3.55 (0.50)*  0.05  4.51 (0.44)*  0.003 | 8.12 (2.13)  7.28 (0.36)  0.54  9.24 (1.94)  0.54 | 9.93 (0.42)  10.02 (0.61)  0.85  9.60 (0.56)  0.46 | 53.50 (4.03)  59.74 (1.45)  0.07  53.50 (6.23)  1.00 | 19.12 (0.84)  14.57 (1.19)*  0.01  17.05 (1.21)  0.07 | 3.00 (0.20)  1.66 (0.94)  0.07  2.30 (2.75)  0.68 | trace  trace  trace |
| **4 N KOH**  wild type  *nks 1-1*  *p-*value  *nks 1-2*  *p-*value | 2.05 (0.27)  2.23 (0.39)  0.56  2.19 (0.02)  0.44 | 0.93 (0.10)  1.10 (0.15)  0.17  1.15 (0.09)  0.05 | 4.88 (1.02)  5.56 (0.47)  0.35  5.93 (0.75)  0.22 | 10.09 (1.21)  10.26 (0.45)  0.84  10.16 (0.61)  0.94 | 55.52 (5.57)  55.73 (0.52)  0.95  53.33 (4.88)  0.63 | 24.43 (2.98)  23.20 (1.57)  0.56  25.39 (2.39)  0.69 | 2.06 (1.23)  1.89 (1.25)  0.88  1.83 (1.17)  0.83 | 0.03 (0.02)  0.03 (0.0)  0.60  0.04 (0.01)  0.35 |

|  | **Fuc**  Avg mol% | **Rha**  Avg mol% | **Ara**  Avg mol% | **Gal**  Avg mol% | **Glc**  Avg mol% | **Xyl/Man**  Avg mol% | **GalA**  Avg mol% | **GlcA**  Avg mol% |
| --- | --- | --- | --- | --- | --- | --- | --- | --- |
| **Total sugars**  wild type  *nks 1-1*  *p-*value  *nks 1-2*  *p-*value | 1.80 (0.14)  1.00 (0.10)*  0.001  2.08 (0.31)  0.22 | 6.19 (0.69)  6.17 (0.51)  0.96  6.13 (0.90)  0.93 | 16.14 (1.10)  17.37 (1.06)  0.24  18.16 (0.49)*  0.04 | 22.91 (1.76)  24.34 (2.51)  0.46  24.26 (1.34)  0.35 | 18.84 (1.81)  21.91 (2.01)  0.12  21.95 (1.84)  0.10 | 13.35 (1.10)  11.81 (0.10)  0.07  11.40 (0.63)  0.06 | 20.78 (2.39)  17.40 (1.25)  0.10  16.02 (0.73)*  0.03 | trace  trace  trace |
