## Supplemental Table 5 for "A DUF1068 protein acts as a pectin biosynthesis scaffold and maintains Golgi morphology and cell adhesion in Arabidopsis"

**Table S5: Additional Golgi and TGN measurements from TEM.** Golgi and TGN from morphometrics as quantified from transmission electron microscopy of high-pressure frozen, freeze-substituted hypocotyl epidermal cells of 3-day-old etiolated seedlings of Col-0 and *nks1-1*  and *nks1-2*. Bold indicates statistically significant differences (p>0.05, t-test for measurements; p>0.01, χ^2^ test for proportions) between wild type and *nks1* mutants.

|  | Col-0 | *nks1-1* | *nks1-2* |
| --- | --- | --- | --- |
| Total number of cells analyzed | 80 | 96 | 75 |
| Total number of Golgi stacks analyzed | 111 | 152 | 112 |
| # Golgi with associated TGN | 79 | 121 | 69 |
| # free TGN observed | 43 | 51 | 31 |
| total # TGN observed | 122 | 172 | 100 |
| % Golgi stacks with associated TGN | 71 | 80 | 61 |
| % TGN associated with a Golgi stack | 65 | 70 | 69 |
| % free TGN | 35 | 30 | 31 |
| Mean # of cisternae per Golgi stack ± S.D. | 4.60 ± 1.15 | **3.92 ± 1.17** | **4.23 ± 1.13** |
| Mean cisternal length ± S.D. (nm) | 1863.99 ± 401.62 | 1843.15 ±424.15 | **1755.67 ± 373.48** |
| Mean cisternal width ± S.D. (nm) | 669.75 ± 213.92 | 788.76 ± 375.20 | 750.24 ± 294.24 |
| Mean cisternal length:width | 2.98 ± 0.89 | 2.81 ± 1.26 | **2.63 ± 0.96** |
