## Supplemental Table 6 for "A DUF1068 protein acts as a pectin biosynthesis scaffold and maintains Golgi morphology and cell adhesion in Arabidopsis"

**Table S6: A list of all Arabidopsis seed lines employed in this study.** uNASC is The European *Arabidopsis* Stock Centre, Nottingham; GABI-KAT is from University of Bielefeld Germany.

| **Plant/Material or Marker line** | **Source of Material** | **Reference** |
| --- | --- | --- |
| *Arabidopsis thaliana* Col-0 | uNASC | (Scholl et al., 2000) |
| *nks1-1* (SALK_151073) | uNASC | (Scholl et al., 2000) |
| *nks1-2* (GK-228H05) | GABI-KAT | (Kleinboelting et al., 2012) |
| *qua2-1* | Gregory Moille | (Mouille et al., 2007) |
| *esmd1-1* | Stephane Verger | (Verger et al., 2016) |
| GFP-HDEL | uNASC | (Batoko et al., 2000) |
| PIN2-GFP | uNASC | (Xu, & Scheres, 2005) |
| Got1p homolog (YFP-WAVE 18) | uNASC | (Geldner et al., 2009) |
| SYP32 (YFP-WAVE 22) | uNASC | (Geldner et al., 2009) |
| VHAa1-GFP | Dr. Karin Schumacher, University of Heidelberg | (Dettmer et al., 2006) |
| RabF2b (ARA7) (RFP-WAVE2) | uNASC | (Geldner et al., 2009) |
| RabF2a (Rha1) (RFP-WAVE7) | uNASC | (Geldner et al., 2009) |
| RabA1e (RFP-WAVE34) | uNASC | (Geldner et al., 2009) |
| NAG-GFP | Dr. Takashi Ueda, University of Tokyo | (Grebe et al., 2003) |
| XYLT-mRFP | Dr. Takashi Ueda, University of Tokyo | (Saint-Jore-Dupas et al 2006) |
| ST-mRFP | Dr. Takashi Ueda, University of Tokyo | (Renna et al., 2005) |
| Sec-n-Rm-2A-Sec GFP | uNASC | (Samalova et al 2006) |
