## Supplemental Table 7 for "A DUF1068 protein acts as a pectin biosynthesis scaffold and maintains Golgi morphology and cell adhesion in Arabidopsis"

**Table S7: A list of all synthetic oligonucleotides used in this study.** Oligonucleotides used in PCR-based experiments were ordered from MWG-Biotech AG (Germany) or Sigma (Australia).

| Gene/T-DNA line | Primer Annotation | Sequence |
| --- | --- | --- |
| NKS1 (At4g30996) | CACC-FP | CACCATGCCTCGTCGATCCGGAGAT |
|  | RP Without STOP | CTCACCTTCCCAACCTGATTGCCGA |
|  | RP with STOP | TTACTCACCTTCCCAACC |
| SALK Insertion primer | LBb1.3 | ATTTTGCCGATTTCGGAAC |
| *nks1-1* (SALK_151073) | LP | GCCTTTTTGGAAAGGAAAAAG |
|  | RP | AGCTCCTCAGTCAGAAGGTCC |
| GABI_Insertion primer | LB | ATATTGACCATCATACTCATTGC |
| *nks1-2* (GK-228H05) | GK-228H05_04nv_LP | TCCAAGTGACTTTCTTGATCCTCT |
|  | GK-228H05_CP70_RP | GGATCTTTCAATCAAAGCTGTCTC |
| NKS1 qRT-PCR primer spanning *nks1-2* | qRT2FP | CAGATGAGCATAGCCGTCATA |
|  | qRT2RP/RT1-RP | CAAAGCTTCAGCTCTCTCTCT |
| NKS1 RT-PCR FP | RT1-FP | GTGCCAACTCTCTTTGTCCT |
| UBI10 (At4g05320) | FP | CACACTTCACTTGGTCTTGCGT |
|  | RP | TAGTCTTTCCGGTGAGAGTCTTCA |
| ACT2 (At3g18780) | FP | AGAGATCACTGCTTTGGCTCC |
|  | RP | ACTCGTCATACTCTGCCTTTG |
| PDF (At1g13320) | FP | TAACGTGGCCAAAATGATGC |
|  | RP | GTTCTCCACAACCGCTTGGT |
| SAND (At2g28390) | FP | AACTCTATGCAGCATTTGATCCACT |
|  | RP | TGATTGCATATCTTTATCGCCATC |
| FADLox (At1g26380) | FP | GACGACACGTAAGAAAGTCC |
|  | RP | CGAACCCTAACAACAAAAAC |
| APT1 (At1g27450) | RT-PCR FP | GAGACATTTTGCGTGGGATT |
|  | RT-PCR FP | CGGGGATTTTAAGTGGAACA |
| BiFC cloning NKS1 (At4g30996) | attB1-FP | GGGGACAAGTTTGTACAAAAAAGCAGGCTTAATGCCTCGTCGATCCGGAGATT |
|  | attB2-RP with STOP | GGGGACCACTTTGTACAAGAAAGCTGGGTATTACTCACCTTCCCAACCTGATTGC |
|  | attB2-RP no STOP | GGGGACCACTTTGTACAAGAAAGCTGGGTACTCACCTTCCCAACCTGATTGCC |
| BiFC cloning QUA1 (At3g25140) | attB1-FP | GGGGACAAGTTTGTACAAAAAAGCAGGCTTAATGGCTAATCACCACCGACTT |
|  | attB2-RP with STOP | GGGGACCACTTTGTACAAGAAAGCTGGGTATCAGAGGCCAAATTGCAAG |
|  | attB2-RP no STOP | GGGGACCACTTTGTACAAGAAAGCTGGGTAGAGGCCAAAATTGCAAGCCT |
| BiFC cloning Got1P  (At3g03180) | attB1-FP | GGGGACAAGTTTGTACAAAAAAGCAGGCTTAATGGCTTCCTTGGAGATGAATG |
|  | attB2-RP with STOP | GGGGACCACTTTGTACAAGAAAGCTGGGTATTAGACAGGGACACGCCTTCCAC |
|  | attB2-RP no STOP | GGGGACCACTTTGTACAAGAAAGCTGGGTAGACAGGGACACGCCTTCCACGGT |
| BiFC cloning IAA1  (At4G14560) | attB1-FP | GGGGACAAGTTTGTACAAAAAAGCAGGCTTAATGGAAGTCACCAATGGGC |
|  | attB2-RP with STOP | GGGGACAACTTTGTATAGAAAAGTTGGGTTAAGGCAGTAGGAGCTTCG |
|  | attB2-RP no STOP | GGGGACAACTTTGTATAGAAAAGTTGGGTGGGCAGTAGGAGCTTCG |
| GO-PROMPTO assay cloning  NKS1 | SfoI-FP | tctggaggaggaGGCATGCCTCGTCGATCCGGAGATT |
|  | KpnI-RP | AGCAGGACTCTAGAGTACTCACCTTCCCAACCTGATT |
| *esmd1-1* dCAPS primers: amplify, then restriction digested by BseXI | FP | GCAGCAAGAGAACGAGGATG |
|  | RP | tgtaaggtttgcaggtgagtg |
| *qua2-1* primers: amplify, then sequence with FP | FP | CCACCTCCCCATGATACTTG |
|  | RP | GAGCTTCCCACTTCAACTGC |
